# Modeling cardiorespiratory coherence in exercise anticipation

**DOI:** 10.1101/2024.03.27.587091

**Authors:** Aditya Koppula, Kousik Sarathy Sridharan, Mohan Raghavan

**Affiliations:** Department of Physiology, Apollo Institute of Medical Sciences and Research, Hyderabad, Telangana, India; Department of Biomedical Engineering, Indian Institute of Technology Hyderabad, Hyderabad, Telangana, India

**Keywords:** Cardiorespiratory Coherence (CRC), Respiratory sinus arrhythmia (RSA), Sino-atrial node (SAN), Hodgkin-Huxley model, Exercise anticipation

## Abstract

Volitional motor activity is associated with a feedforward cardiorespiratory response to actual or impending movements. We have previously shown in the CRC study that the expectation of physical exercise causes a decrease in cardiorespiratory coherence that scales with the anticipated load. The present work uses a modeling approach to investigate the mechanisms that can cause a fall in cardiorespiratory coherence (CRC). We devised a Hodgkin-Huxley model of a cardiac pacemaker cell using the NEURON module. We simulated the effect of autonomic tone, sympathetic & respiratory-vagal modulation, and respiratory irregularity on pacemaker cell output by injecting efflux/influx current to model the parasympathetic/sympathetic effects, respectively. The vago-sympathetic tone was modeled by altering the direct current bias of the injected current and the respiratory-vagal effect by the periodic modulation of the injected current at a frequency of 0.2 Hz, corresponding to a respiratory rate of 12 breaths/min. Sympathetic modulation was simulated by injecting a low-frequency current close to Mayer wave frequency (0.08 Hz). We computed the coherence between the instantaneous pacemaker rate and respiratory-vagal modulation current as a model analog to experimental CRC. We found that sympathetic modulation, low vagal tone/high sympathetic tone, and respiratory irregularity can cause a decrease in CRC. We corroborated the model results with the actual data from the CRC study. In conclusion, we employ a novel approach combining insights from the experimental study and a physiologically plausible modeling framework to understand the mechanisms underlying the fall of cardiorespiratory coherence induced by the expectation of exercise.

**NEW & NOTEWORTHY**

Cardiorespiratory coherence is diminished in response to respiratory irregularity, low vagal/high sympathetic tone, and prominent low-frequency sympathetic modulation.

Expectation of physical activity induces respiratory irregularity and increased sigh frequency and that contributes to diminished cardiorespiratory coherence in expectation of exercise.

There is a greater fall of coherence with the non-linear (logistic) transformation of injected current, indicating the non-linear nature of cardiorespiratory interactions preceding the onset of exercise.

## INTRODUCTION

Living organisms anticipate and evoke predictive or feedforward responses to the challenges or demands imposed on them. The challenge of physical exercise requires multiorgan system adjustments, including heightened cardiac and respiratory outputs in parallel with actual or expected motor output. Several studies have explored the cardiac or respiratory responses to actual or anticipated physical exercise(1–7), while we focus on the coordinated cardiorespiratory activity, also termed cardiorespiratory coupling. Cardiorespiratory coupling encompasses physiologically distinct phenomena like cardiorespiratory phase synchronization and respiratory sinus arrhythmia (RSA)(8,9). RSA is the rise and fall of heart rate coinciding with the phases of respiration (10) and can be quantified by “Coherence” between heart rate and respiratory signal. We investigate the changes induced in coherence by the expectation of physical exercise. In a previously published study (**CRC study**)(11), we have shown that cardiorespiratory coherence (CRC) is decreased in expectation of exercise in a load-dependent manner (Appendix **Figures A1-A3**). In the CRC study, young adult participants were subjected to different types and intensities of isometric & isotonic exercises. The ECG-derived heart rate and the abdominal belt plethysmography-derived respiratory signal, recorded in the 5 minutes of anticipation phase before the onset of exercise, were used to compute the coherence. We briefly summarize the experimental design in the Appendix (Appendix **Figure A1**). Coherence is an abstract frequency domain metric, and it is far from obvious what the change in coherence means in terms of cardiac, respiratory, and autonomic physiology. An insight into the underlying mechanisms can help decipher the causes & consequences of a fall in cardiorespiratory coherence and positively affect its interpretability & usability. This work attempts to answer two closely related but distinct questions using a modeling approach: 1. What physiological factors affect coherence between heart rate and respiration? 2. What are the mechanisms that can explain the decrease in cardiorespiratory coherence with the expectation of exercise? We demonstrate a hybrid paradigm using experimental and modeling insights to uncover the underlying physiological mechanisms of coherence fall.

As coherence quantifies the efficacy of heart rate modulation by respiration, we chose a reductionist approach based on a model of a pacemaker cell and the mechanisms that modulate its spike rate. The primary pacemaker of the human heart is the Sino-trial node (SAN), while the modulation of pacemaker behavior is mediated by the efferents from the two arms of the autonomic nervous system, viz., Sympathetic and parasympathetic nervous system (vagus). The modulatory nerve impulse traffic in the vagus and sympathetic nerve fibers innervating the SAN can be conceptually dissected into two components: 1. “Tonic component” or “DC bias” represented by the average spike frequency 2. “Phasic component” is the quasi-periodic fluctuation in firing rate/spike frequency around the DC-bias(12–16). In the vagus, phasic modulation in spike frequency is induced mainly by the signal that “spills over” from the respiratory central pattern generator (CPG) in the brain stem(10). Therefore, respiratory-vagal activity induces SAN spike frequency variation at the respiratory frequency(0.15-0.4 Hz). In contrast, sympathetic modulation is thought to occur at low frequencies (< 0.15 Hz), partly due to the slow transmitter kinetics of sympathetic neurotransmitters (17,18). The most prominent low-frequency trend attributed to the sympathetic modulation is the “Meyer waves” (0.06-0.1 Hz)(19). We have two-fold objectives for modeling: 1. Develop a single-cell model of the cardiac pacemaker cell and demonstrate pacemaker rate modulation in response to respiratory-autonomic input 2. To explore the underlying mechanisms for the fall in pre-exercise coherence.

## MATERIALS AND METHODS

We simulated the cardiorespiratory coherence at rest and the pre-exercise state as follows: 1. A model of the pacemaker cell of the heart is devised 2. The autonomic and respiratory modulatory mechanisms on the pacemaker cell are simulated 3. We perform preliminary “sanity-check” experiments on the pacemaker model to assess its response to a subset of autonomic-respiratory modulatory mechanisms, including autonomic tone, vagal & sympathetic modulation, and respiratory patterns 4. “Full space” exploration of the effect of all combinations of autonomic-respiratory factors/mechanisms is done to uncover the mechanisms of anticipatory fall in cardiorespiratory coherence.

### Model of pacemaker cell

A Hodgkin-Huxley-type cell model of a cardiac pacemaker was devised using ***NEURON***(20) module in Python software(20). A parsimonious set of relevant ion channels was inserted into the model cell membrane (**Table 1**). The model cell has a length of 100 μm, a diameter of 20 μm, and a membrane capacitance of 2 μF/cm^2^. The model does not incorporate sarcoplasmic reticulum(SR)-based calcium storage/ release/ reuptake mechanisms. The model cell and ion channel parameters were iteratively adjusted to get a spontaneous pacemaker rate close to the intrinsic frequency of the human SAN cell in a completely denervated-isolated heart (21,22).

**Table 1:**
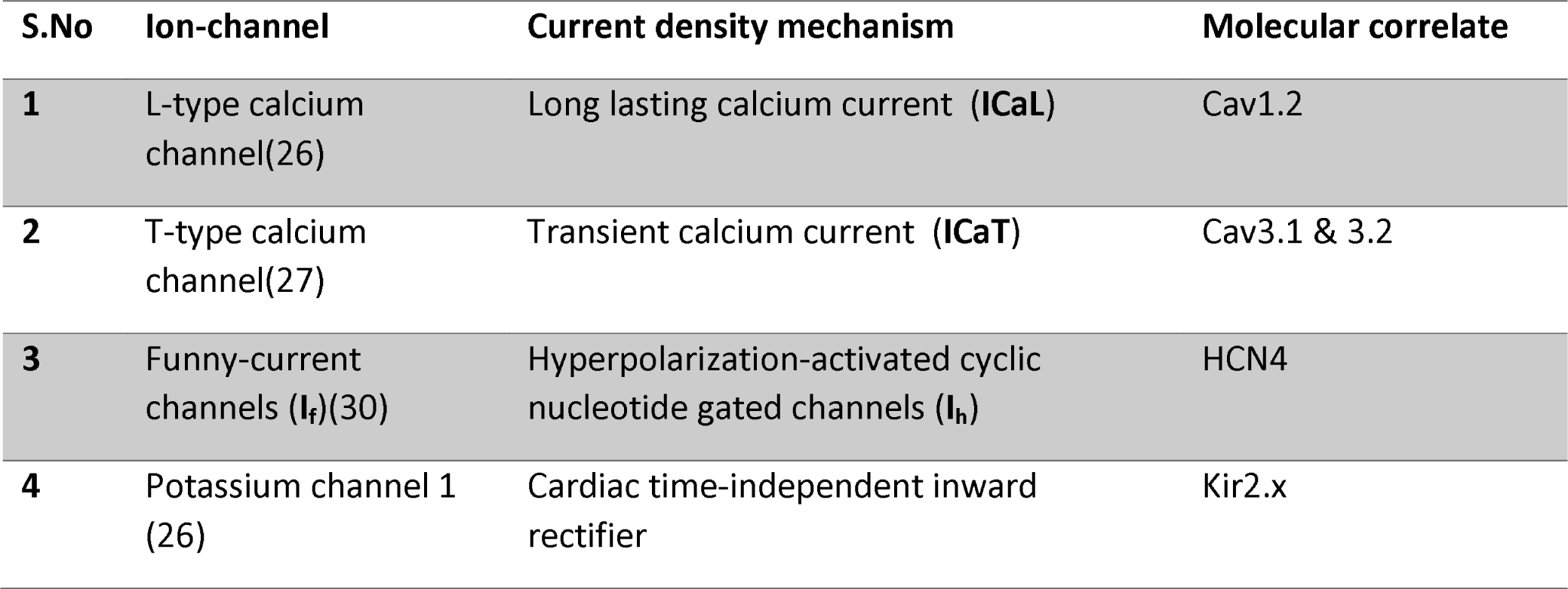
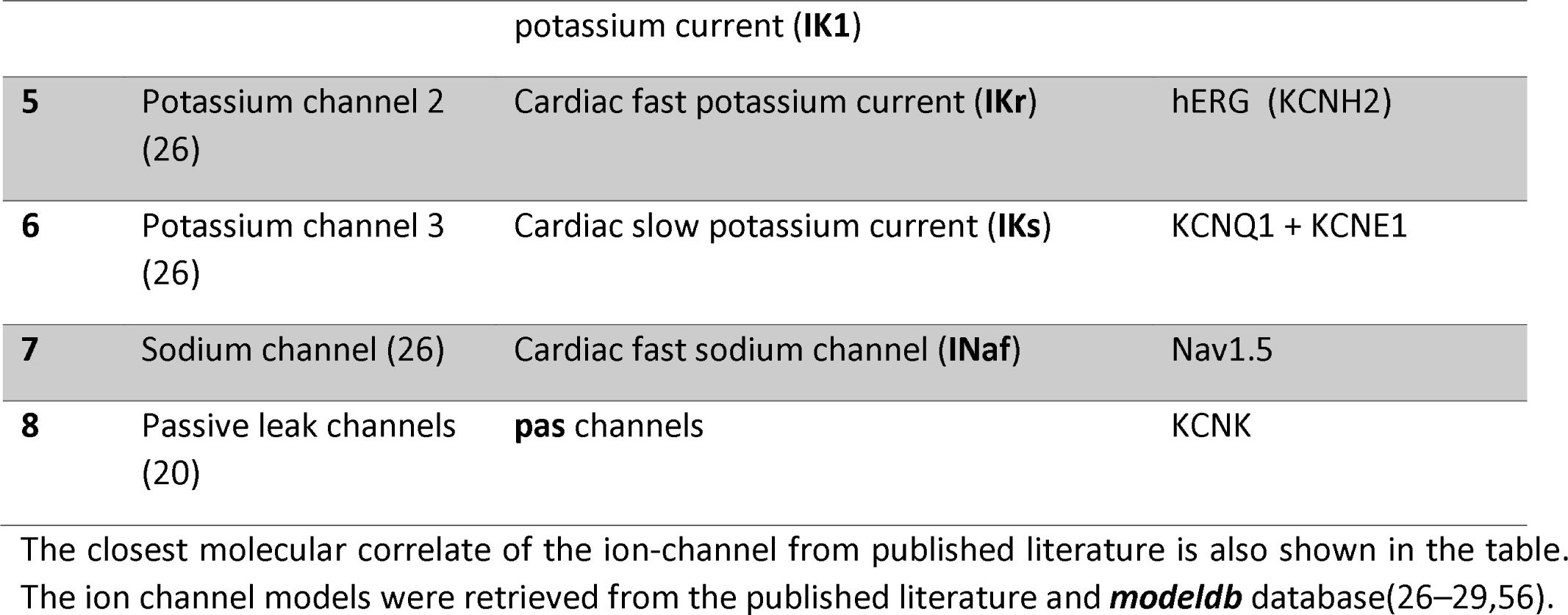
Ion-channels used in the model of cardiac pacemaker cell.

### Simulation of autonomic and respiratory modulatory mechanisms

The effect of autonomic modulation on the pacemaker was simulated by injecting current into the model cell using a current clamp point-process mechanism using the NEURON module. The current clamp was switched on 20 seconds after the onset of the simulation to allow for the stabilization of the pacemaker rate. In health, the vagus increases potassium efflux out of the cell, while the sympathetic system increases sodium/calcium influx into the cell(23–25). Our model simulated vagal activity by injecting a negative valued current into the cell (efflux current). In contrast, sympathetic activity was simulated by injecting a positive valued current (influx current). We simulated the effect of tonic and phasic components of the autonomic nervous system effect on the SAN by changing the pattern of the injected current. The injected current comprises of the following components:

1. **A constant or Direct current (DC) component**: The DC component is a zero-to negative-valued constant current that determines the baseline bias of the injected current. A more negative valued DC bias implies a high vagal tone and/or low sympathetic tone, while a less negative to zero DC bias indicates a low vagal tone and/or high sympathetic tone. We tested the effect of several values of DC bias to assess the effect of vago-sympathetic tone.
2. **High**-**frequency component**: A phasic current with a frequency ∼0.2 Hz (corresponding to a breath rate of 12/minute) was injected as the high-frequency component into the model cell to simulate the effect of respiratory-vagal modulation. It is well known that respiration exerts its action on the pacemaker rate through the vagus. We evaluated the effect of two patterns of breathing:

a. Sinusoidal breathing at 0.2 Hz
b. **Respiratory irregularity:** Natural breathing has variability in rate & depth, and is sporadically punctuated by augmented deep breaths called “sighs”. Therefore, we simulated the effect of respiratory rate variability (**rrv**), respiratory amplitude/depth variability (**rav**) and **sighs**. The simulation pipeline for respiratory irregularity is described in the **Appendix**. The waveforms of the simulated sinusoidal and variable/irregular breathing patterns are shown in **Figure 1**.
3. **Low-frequency component**: A sinusoidal current injection at a frequency of 0.08 Hz was used to represent the “sympathetic modulation” effect. Heart rate variability (HRV) literature indicates that sympathetic modulation occurs in the low-frequency band (0.04-0.15 Hz). The Mayer waves in blood pressure (and heart rate), the well-known consequence of sympathetic and baroreflex modulation of SA node activity, peak in the frequency band of 0.06-0.1 Hz(19).

**Figure 1:**
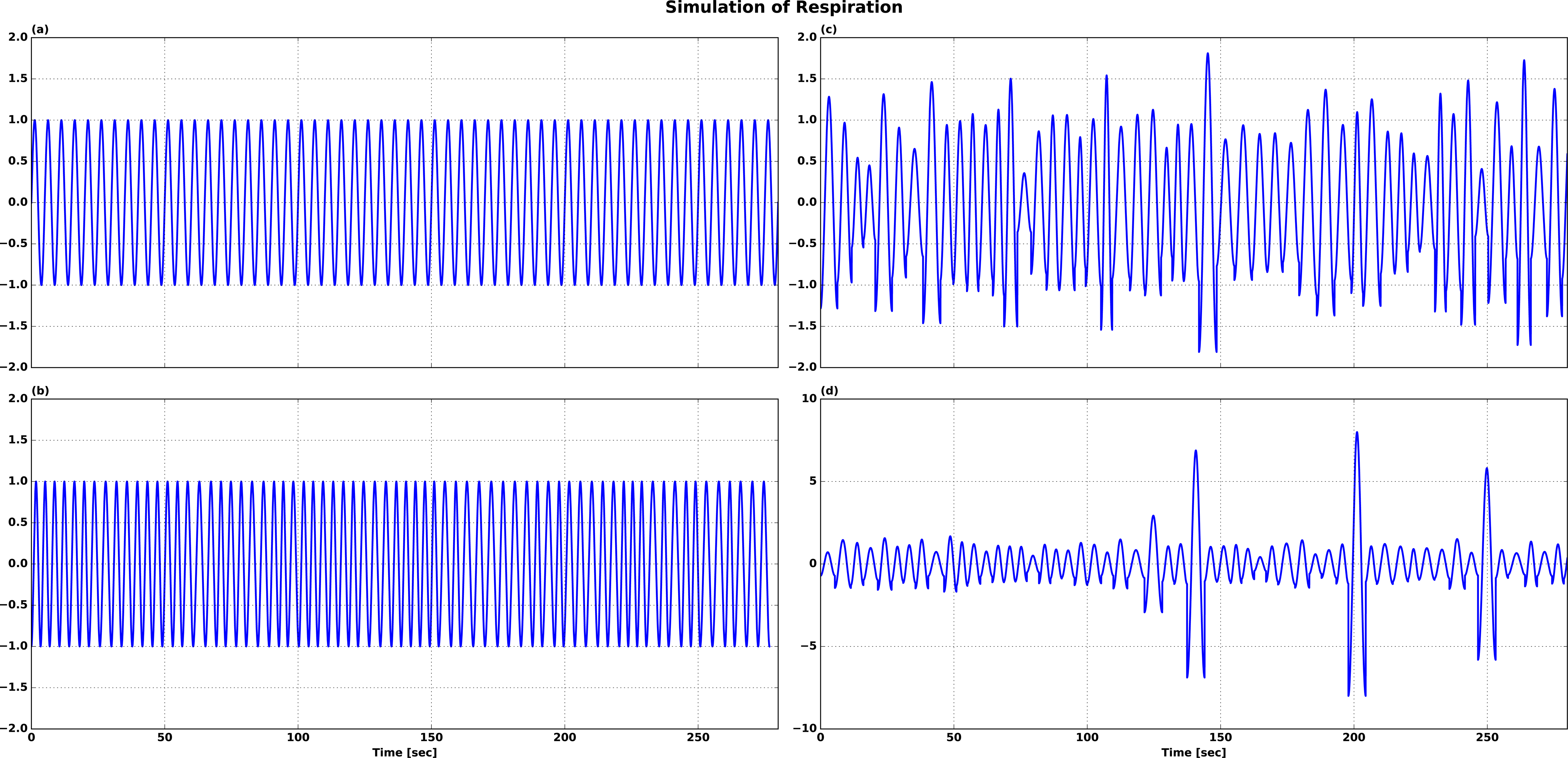
Simulation of Physiological breathing patterns. (A)-This shows a sinusoidal breath pattern at 0.2 Hz. (B)-Respiratory frequency varies from breath to breath between 0.15-0.25 Hz (RRV: respiratory rate variability). (C)-Respiratory rate and amplitude variability (RRAV). (D)-Intermittent “sighs” in the background of respiratory rate and depth variability (RRAVS).

All the simulations were performed for 320 seconds at a sampling period of 0.025 milliseconds. The first 20 seconds allowed the stabilization of the pacemaker rate, and the current injection corresponding to the respiratory-autonomic modulation was confined to the subsequent 300 seconds, which is consistent with the experimental design in the CRC study(11). The simulated respiration (High-frequency component) and/or sympathetic modulation signal (low-frequency component) were injected as a current into the model cell, and the effects on pacemaker rate fluctuations and model ***cardiorespiratory coherence*** (coherence between instantaneous pacemaker rate and the respiratory signal i.e., the High-frequency component) were studied. We performed the simulations under two conditions: 1. The respiratory/autonomic modulation current was injected without any transformation. 2. The respiratory/autonomic modulation current was transformed with a logistic function (**Equation 1**) before the current injection. We show that coherence begins to fall only if the input current is transformed by the logistic function (before injection into the cell) and respiratory-vagal modulation is variable/irregular.

**Equation 1: A 2-parameter logistic function**. The input data is x (injected current) and the parameter K is computed from the 75th percentile & interquartile range (IQR) of the input data (K=75th percentile value + 5×IQR). The x_0_ is the median of the input data and b is set to be 1. The median and IQR-based parameters for the logistic function make the transform robust to the “sigh” like outliers in the input

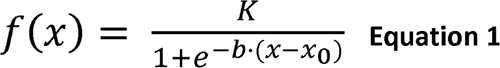

### Sanity-check experiments

The objective of sanity-check experiments is to reproduce a general class of results that are characteristic of normal SAN cell behavior and determine the conditions/mechanisms that can cause a fall in coherence. We evaluated the effect of a select subset of mechanisms on pacemaker output, including the following: **1.** Autonomic tone (DC bias) **2.** Sinusoidal breathing superimposed on different levels of autonomic tone with/without logistic transformation **3.** Respiratory variability: rate & depth variability, sighs, and sympathetic modulation with and without logistic transformation. We assess the following in pacemaker model output: membrane potential, injected current, instantaneous pacemaker rate, and magnitude-squared coherence (between instantaneous pacemaker rate and the high-frequency component of injected current, which is the surrogate for respiratory-vagal signal). The magnitude-squared coherence is computed as shown in **Equation 2**. Both time series (pacemaker rate & high-frequency component) were down-sampled to 4 Hz, and coherence was computed using scipy.signals.coherence() function from the **python** scipy package with the following settings: Welch’s method, nfft=256, overlap=128, and mean detrending.

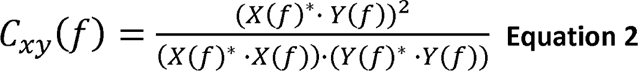

**Equation 2: Magnitude-squared coherence.** If *x* and *y* are two signals (like pacemaker rate and respiration/high-frequency signal), *Cxy(f) =* Coherence between *x* and *y* as a function of frequency, *f* = frequency, *X(f)* = FFT (fast fourier transform) of time series *x*, *X(f)\** = complex conjugate of *X(f), Y(f)* = FFT of time series *y, Y(f)* =* complex conjugate of *Y(f)*.

In the following, we outline the simulation pipeline for the effect of autonomic tone, sinusoidal breathing, and respiratory variability:

1. Autonomic tone: The effect of steady autonomic tone was evaluated at several values of the current clamp including 0 nA, −1 nA, −1.8 nA, and −2.5 nA, corresponding to increasing vagal tone/decreasing sympathetic tone with more negative DC bias.
2. Sinusoidal breathing: We simulated respiratory modulation of vagal activity by injecting a high-frequency sinusoidal current at a frequency of 0.2 Hz (and an amplitude of 0.3 nA) centered around three different magnitudes DC bias/vagal tone (−0.3 nA, −1 nA, −2 nA), to demonstrate the role of vagal tone on the modulation-induced pacemaker rate variability. As the vagal tone determines the average heart rate, the above-mentioned simulation is analogous to the effect of average heart rate on heart rate variability (HRV) in general and RSA in particular. We tested the interaction between the amplitude of respiratory-vagal modulation (sinusoid amplitude=0.3 nA, 0.6 nA), and steady vagal/sympathetic tone (DC bias= −0.6 nA, −2 nA). The effect of the amplitude of respiratory-vagal modulation corresponds to that of volume/depth of breathing, which is the volume of the air inspired or expired at rest in health. We also assessed cardiorespiratory coherence with factorial combinations of respiratory/vagal modulation and autonomic tones. We show that peak coherence is close 1 with any combination of sinusoidal respiratory/vagal input and untransformed/transformed current injection.
3. Respiratory variability: We evaluated the effect of respiratory rate & depth variability and sighs on the model pacemaker output and coherence. A constant negative current (−1.5 nA) was added to the respiratory signal before injection into the pacemaker cell to account for vagal tone. Sympathetic modulation was modeled by injecting a low-frequency sinusoid at 0.08 Hz. The following combinations were assessed: 1. respiratory rate variability 2. respiratory rate & depth variability 3. respiratory rate & depth variability with sighs 4. respiratory rate & depth variability, sighs and sympathetic modulation with different levels of autonomic tone. The procedure and profile of various types of simulated variabilities are presented in supplementary information. We demonstrate that coherence decreases only after introducing respiratory variability and logistic transformation of injected current into the model.

### Full-space exploration of mechanisms for expectation-induced coherence fall

The objective of “full-space” exploration is to uncover the possible mechanisms responsible for the expectation-induced fall in coherence in the CRC study(11). We performed a 2-step simulation and analysis: **Step 1.** We simulated the effect of all the possible combinations of the factors/conditions on the pacemaker, including autonomic tone, sympathovagal modulation, and respiratory irregularity. About 96 combinations of factors/mechanisms (8×12=96; 8 in **Table 2** & 12 in **Table 3**) were simulated for 20 iterations in each combination. **Step 2.** For each of the 11 experimental conditions in the published CRC study(11), three best fits from the 96 simulated ones were retrieved to understand the possible mechanisms behind the fall in cardiorespiratory coherence preceding exercise.

**Table 2:** Set 1. Factors pertaining to autonomic tone and sympathetic modulation.

| S.No | Factor/condition | Features |
| --- | --- | --- |
| **1** | AT | injected current has zero-DC bias; interpreted as the lowest level of vagal tone and/or highest sympathetic tone, no sympathetic modulation. |
| **2** | LT | injected current has a negative DC-bias of (-1) nA; interpreted as the low level of vagal tone and/or high sympathetic tone, no sympathetic modulation. |
| **3** | BT | injected current has a negative DC-bias of (-1.5) nA; interpreted as the moderate level of vagal tone and/or moderate-to-low sympathetic tone, no sympathetic modulation. |
| **4** | HT | injected current has a negative DC-bias of (-2) nA; interpreted as the highest level of vagal tone and/or lowest sympathetic tone, no sympathetic modulation. |
| **5** | SM_AT | presence of sympathetic modulation (0.08 Hz low-frequency component); zero-DC bias; lowest vagal tone/highest sympathetic tone. |
| **6** | SM_LT | presence of sympathetic modulation (0.08 Hz low-frequency component); negative-DC bias of (-1) nA; low vagal tone/high sympathetic tone. |
| **7** | SM_BT | presence of sympathetic modulation (0.08 Hz low-frequency component); negative-DC bias of (-1.5) nA; moderate vagal tone/moderate-to-low sympathetic tone. |
| 8 | SM_HT | presence of sympathetic modulation (0.08 Hz low-frequency component); negative-DC bias of (-2) nA; highest vagal tone/lowest sympathetic tone. |

**AT: Absent/No Tonic current injection; LT: Low Tonic current injection; BT: Basal Tonic current injection; HT: High Tonic current injection.** AT, LT, BT, HT are terms indicating DC bias in current clamp of the pacemaker model cell to simulate vagal/sympathetic tone. The factors/conditions 1-4 & 5-8 above are differentiated by the absence ( conditions 1-4) or presence (conditions 5-8) of sympathetic modulation respectively. AT → HT is characterized by increasing vagal tone or decreasing sympathetic tone.

**Table 3:** Set 2. Factors pertaining to vagal modulation/ respiratory variability.

| S.No | Factor/condition | Features |
| --- | --- | --- |
| 1 | Sinusoidal | respiratory signal is sinusoidal at 0.2 Hz (no rate/depth variability or sighs). |
| 2 | rrv1 | respiratory rate variability pattern 1 (power law exponent=0.3, mean frequency=0.2 Hz); no amplitude variability or sighs. |
| 3 | rrv2 | respiratory rate variability pattern 2 (power law exponent=0.8, mean frequency=0.265 Hz); no amplitude variability or sighs |
| 4 | rav | respiratory amplitude variability at a constant frequency of 0.2 Hz; no rate variability or sighs. |
| 5 | s | Sighs sporadically interspersed among sinusoidal breaths; except for sighs, no rate & depth variability. |
| 6 | rrav1 | respiratory rate variability pattern 1 + depth variability; no sighs |
| 7 | rrav2 | respiratory rate variability pattern 2 + depth variability; no sighs |
| 8 | rrvs1 | respiratory rate variability pattern 1 + sighs; no amplitude variability |
| 9 | rrvs2 | respiratory rate variability pattern 2 + sighs; no amplitude variability. |
| 10 | ravs | respiratory amplitude variability + sighs for the signal at 0.2 Hz frequency; no rate variability. |
| 11 | rravs1 | respiratory rate variability pattern 1 + depth variability+ sighs |
| 12 | rravs2 | respiratory rate variability pattern 2 + depth variability+ sighs |

Factors/Conditions 1-12 represent different patterns of respiratory variability

Of the three principal coherence modifying factors (viz., respiratory variability, autonomic tone, sympathovagal modulation) assessed in **step 1** above, the role of respiratory variability can be corroborated by simple comparison with the respiratory variability in the rest/anticipatory phase of the experimental data in the CRC study. In contrast, the role of autonomic tone/sympathovagal modulation can only be indirectly inferred from the best-model fit approach in **step 2.** The CRC study (11) consisted of the following exercises: **(i)**. baseline **(ii)**. hand-grips (4 types): low-intensity handgrip of the left hand, low-intensity handgrip of the right hand, high-intensity handgrip of the left hand, high-intensity handgrip of the right hand. **(iii)**. Bicycle exercise (6 types): left-sided low intensity, right-sided low intensity, left-sided high intensity, right-sided high intensity, bilateral low intensity, bilateral high intensity. The best fits were determined based on the mean squared error (mse) between the **average ‘model’ coherence** (from each of the 96 combinations) and the **average ‘experimental’ coherence** for 11 exercise conditions. All the simulations in “full space” exploration involved the injection of currents transformed by the logistic function.

## RESULTS

The original and the modified ion-channel parameters of the pacemaker cell model are shown in **Table 4** (20,26–31). The pacemaker cell fires spontaneously at a steady rate of 105 spikes/min (**Figure 2 & Figure 3**), corresponding to the state of an isolated SAN that is devoid of any extrinsic neuro-humoral influence(22,32,33).

**Figure 2:**
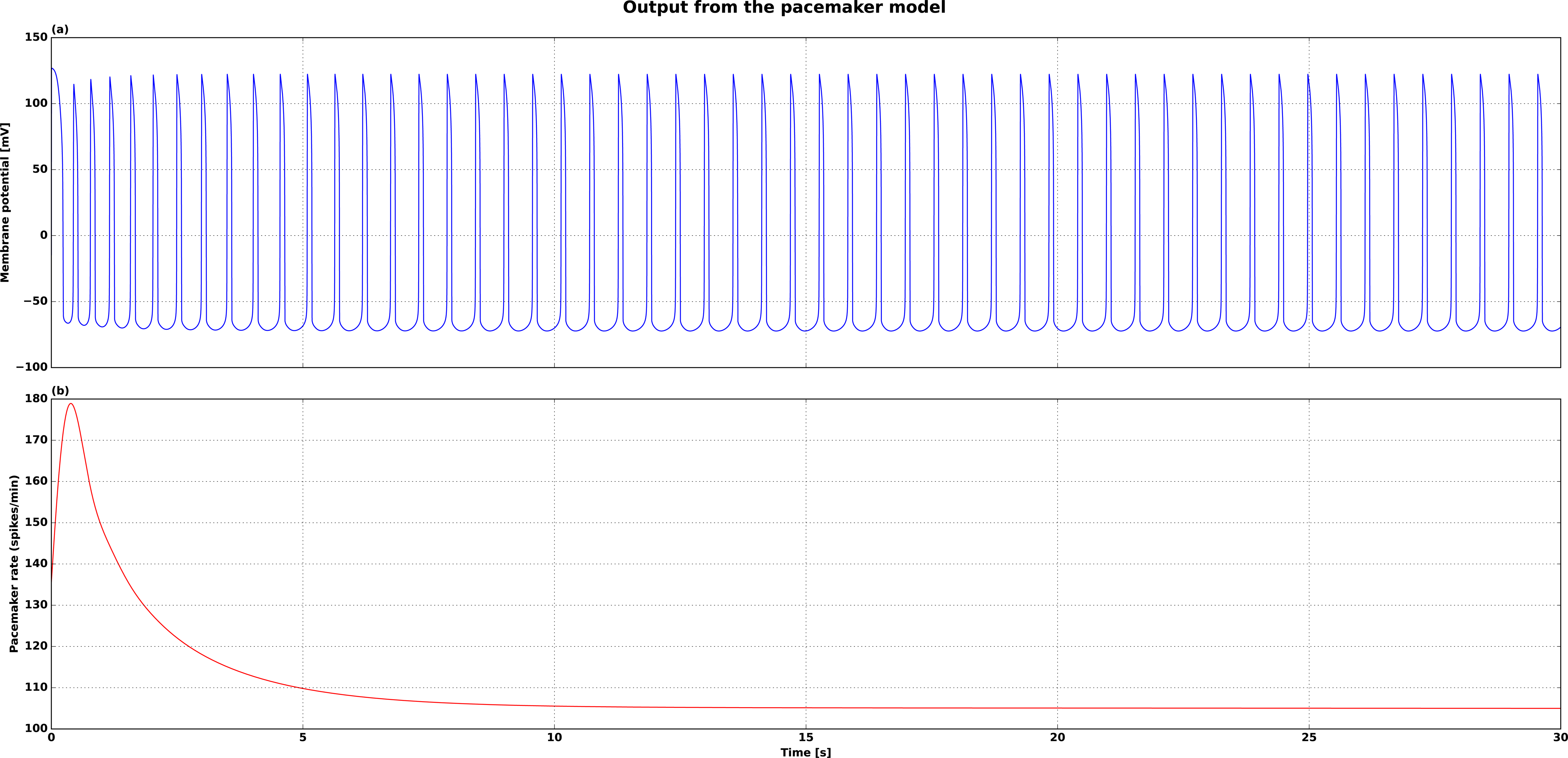
Pacemaker model cell spontaneous output. After the onset of simulation, the spontaneous output of the pacemaker model cell takes ∼ 10 seconds to attain a steady state firing rate of 105 spikes/min. No current was injected into the cell in this simulation. In the subsequent simulations, current was injected at 20 seconds after simulation onset that allowed stable pacemaker before current injection At the steady state, there is no appreciable heart rate variability. This model corresponds to the state of a normal SA node that is completely isolated from all the external influences.

**Figure 3:**
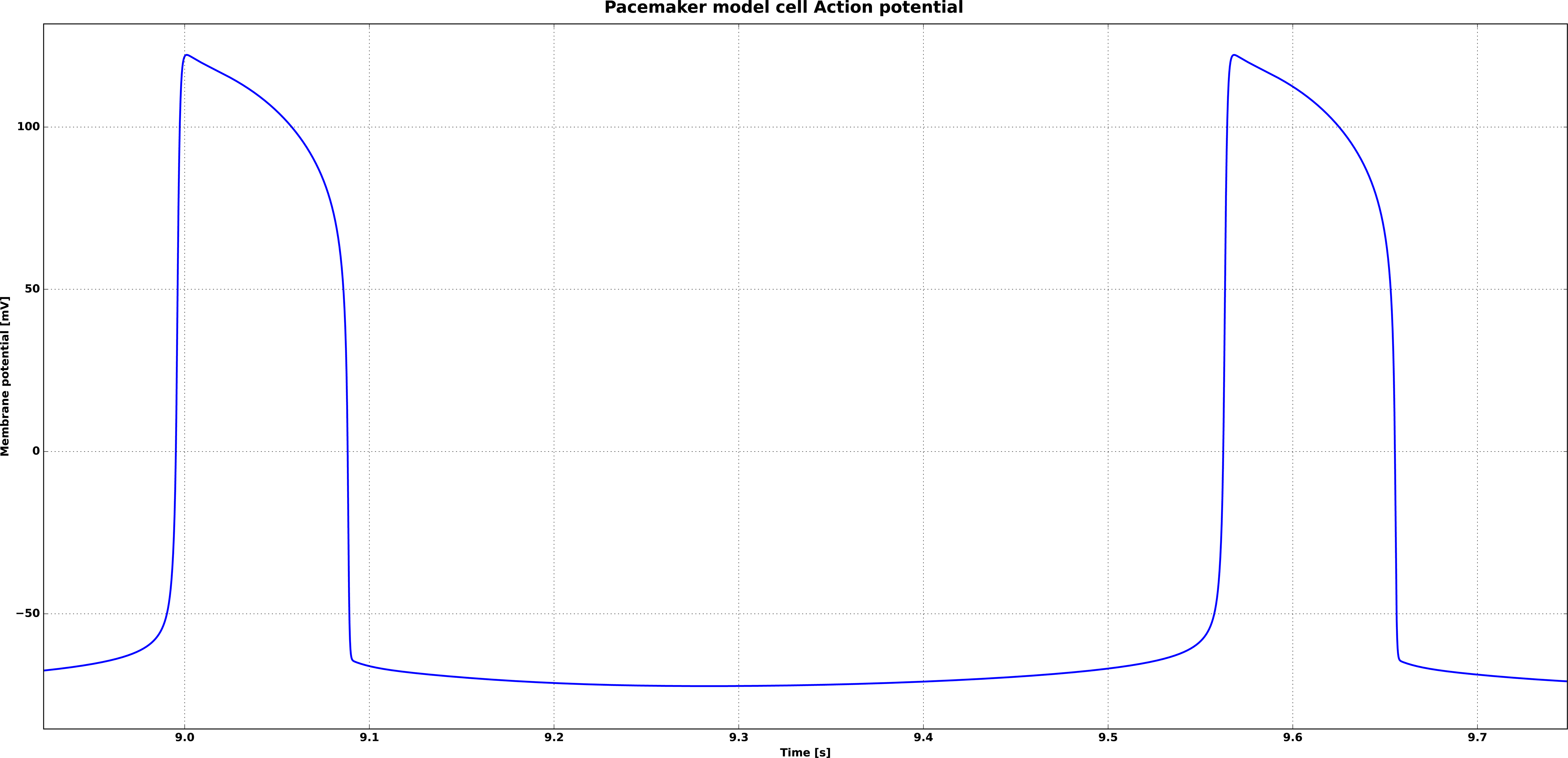
Magnified view of pacemaker model cell action potential. The various phases of the model cell action potential, including the plateau and the spontaneous ‘diastolic’ depolarization phases, are shown.

**Table 4:** The ion channels and the parameters modified for the pacemaker cell model.

| S.No | Ion channel | Original parameters | Modified parameters |
| --- | --- | --- | --- |
| 1 | ICaT(27) | $\overline{g_{CaT}} = 0.008 \text{ mho. cm}^{-2}$ | $\overline{g_{CaT}} = 0.094 \text{ mho. cm}^{-2}$ |
| 2 | I<sub>h</sub>(30) | $\overline{g_h} = 0.001 \text{ S. cm}^{-2}$<br>$E_h = 0 \text{ mV}$ | $\overline{g_h} = 1e - 5 \text{ S. cm}^{-2}$<br>$E_h = -10 \text{ mV}$ |
| 3 | IK1(26) | $\overline{g_{K1}} = 0.0001567 \text{ S. cm}^{-2}$ | $\overline{g_{K1}} = 0.0005567 \text{ S. cm}^{-2}$ |
| 4 | IKr(26) | $\overline{g_{Kr}} = 0.0000588 \text{ S. cm}^{-2}$<br>$\tau_{act} = 1 \text{ ms}$ | $\overline{g_{Kr}} = 0.01 \text{ S. cm}^{-2}$<br>$\tau_{act} = 1.2 \text{ ms}$ |
| 5 | IKs(26) | $\overline{g_{Ks}} = 0.0000588 \text{ S. cm}^{-2}$ | $\overline{g_{Ks}} = 0.000588 \text{ S. cm}^{-2}$ |
| 6 | INaf(26) | $\overline{g_{Naf}} = 0.0156 \text{ S. cm}^{-2}$<br>$\tau_{act} = 1 \text{ ms}$ | $\overline{g_{Naf}} = 0.0008 \text{ S. cm}^{-2}$<br>$\tau_{act} = 0.2 \text{ ms}$ |
| 7 | Pas(20) | $\overline{g_{pas}} = 0.001 \text{ S. cm}^{-2}$<br>$E_{pas} = -70 \text{ mV}$ | $\overline{g_{pas}} = 1e - 6 \text{ S. cm}^{-2}$<br>$E_{pas} = -80 \text{ mV}$ |

Following the modification of the ion channel parameters, the pacemaker rate is 105 beats/min. $\overline{g_{\{x\}}}$ is the maximum conductance and $\overline{E_{\{x\}}}$ is the reversal potential of the channel $x$ . $\tau_{act}$ is the activation time constant of the channel.

### Sanity-check results

#### Autonomic tone and Sinusoidal breathing

In what follows, we discuss the results of simulating vagal tone and sinusoidal respiratory-vagal modulation. Injection of constant negative current (corresponding to steady vagal tone) resulted in a decline of pacemaker rate, with lower pacemaker rate at stronger vagal tone (more negative current injection) (**Figure 4**). In can also be seen that there is no heart rate variability with steady vagal tone. The effect of periodic respiratory vagal modulation on heart rate was simulated with a sinusoidal current injection in the model cell. The sinusoidal current injection at any DC bias showed periodic fluctuations in heart rate at the same frequency, mimicking the RSA. We show that the amplitude of RSA (pacemaker rate fluctuations at respiratory frequency) was greater with a stronger vagal tone (**Figure 5 & Table**).**5** The effect of depth of breathing was simulated by changing the amplitude of injected sinusoid, and we show that larger the amplitude of sinusoid (deeper the breathing), greater the magnitude RSA fluctuations and vice versa (**Figure 6 & Table**)**6**. For a given sinusoid amplitude, the magnitude of RSA fluctuation was greater with stronger vagal tone indicating interaction between depth of breathing and level of tone. The sanity-check results presented so far are consistent with the behavior of SAN in healthy humans supporting the validity of model results. Coherence between the sinusoidal “respiratory frequency” modulation current and the instantaneous pacemaker rate shows a peak value of 1 around the modulation frequency of 0.2 Hz, and the coherence did not differ between various levels of vagal tone and modulation (**Figure 7 (A)**). Coherence results indicate that sinusoidal breathing with or without logistic transformation cannot account for expectation induced fall in cardiorespiratory coherence (**Figure 7 (A) & (B)**).

**Figure 4:**
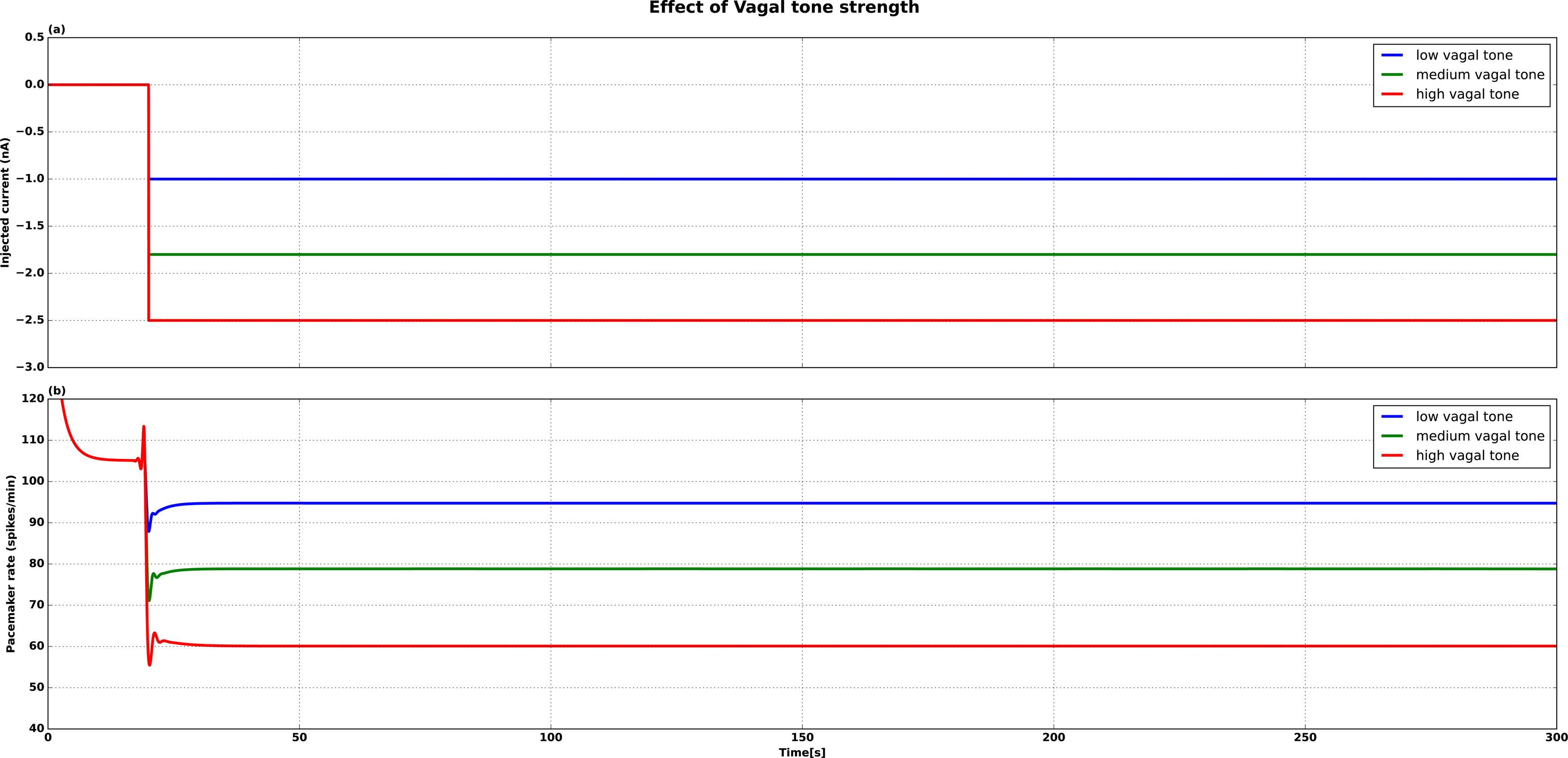
Effect of vagal tone. Vagal tone was increased by increasing the injected current (low: −1 nA, medium: −1.8 nA & high vagal tone:-2.5 nA). With increasing vagal tone, pacemaker rate drops precipitously.

**Figure 5:**
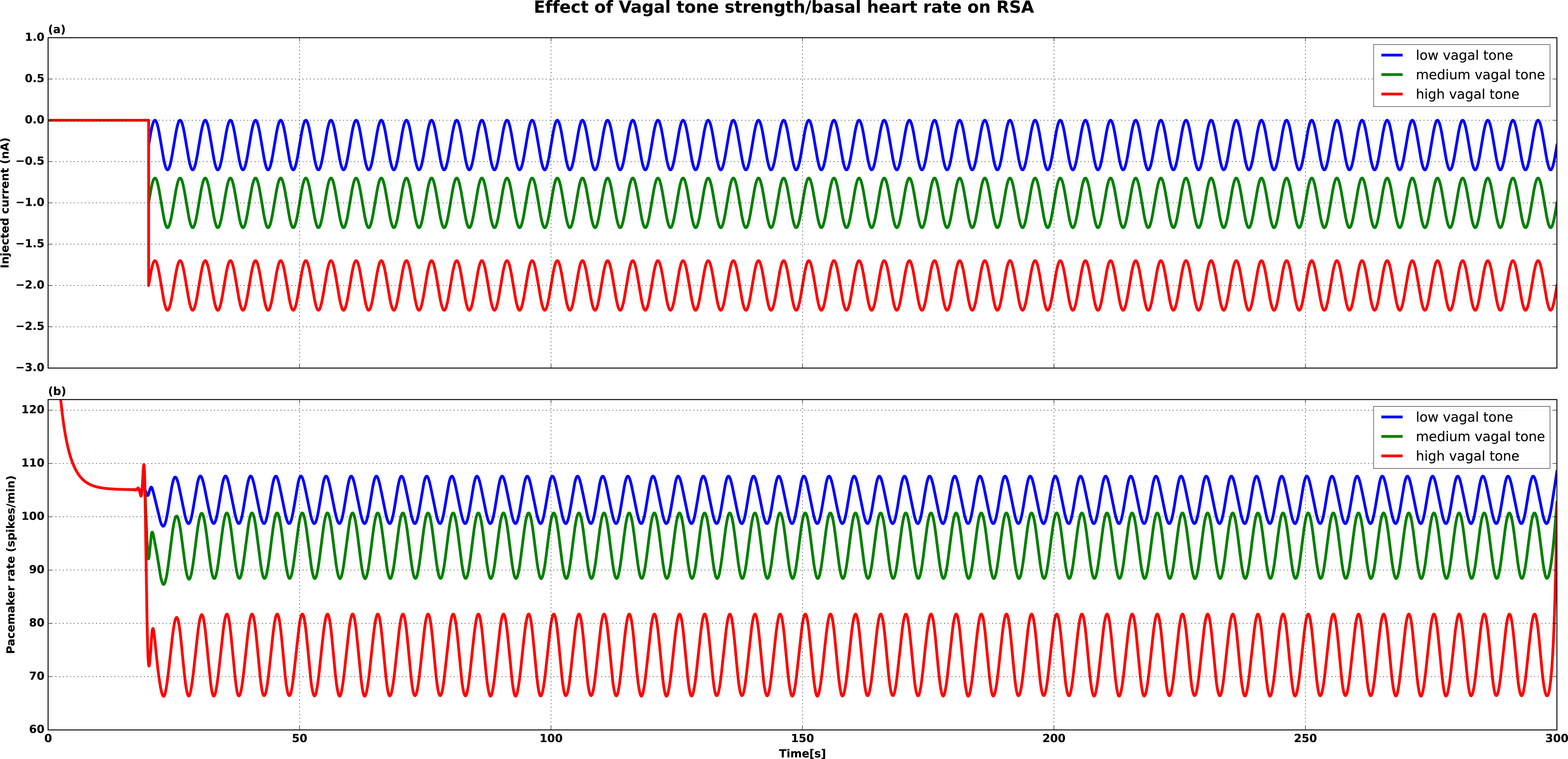
Effect of vagal tone on RSA-like pacemaker rate fluctuations. For the identical respiratory-vagal modulation (amplitude = 0.3 nA, frequency = 0.2 Hz), higher the vagal tone (Magnitude of injected steady current: low vagal tone = −0.3 nA, medium vagal tone = −1 nA, High vagal tone = −2 nA), lower the average pacemaker rate and greater the amplitude of pacemaker rate variations.

**Figure 6:**
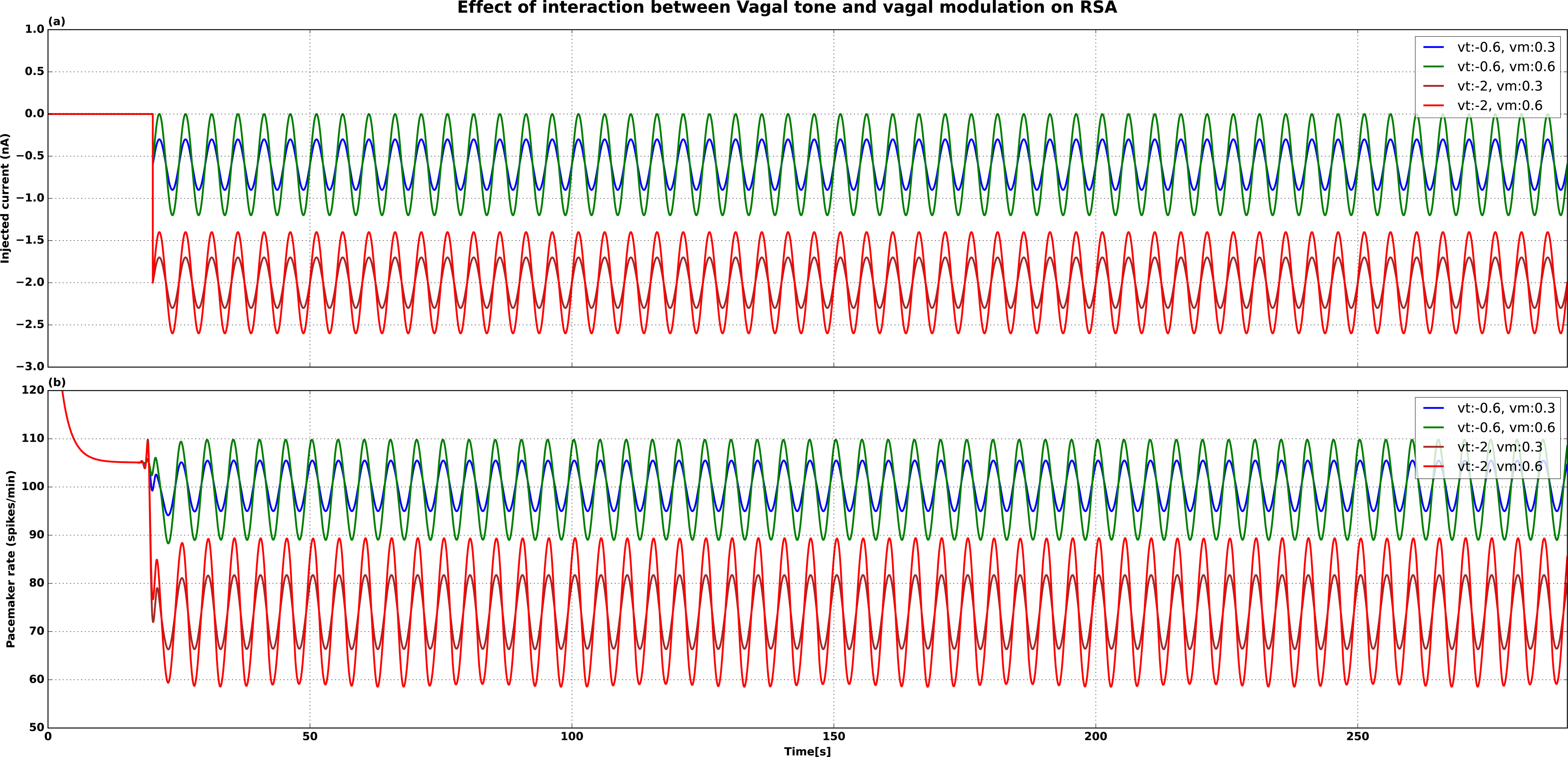
Effect of Vagal tone on respiratory-vagal modulation of pacemaker rate. vt: vagal tone (low vagal tone =-0.6 nA, High vagal tone= −2 nA), vm: vagal modulation (low vagal modulation = 0.3, high vagal modulation = 0.6). Doubling the strength/amplitude of vagal modulation from 0.3 to 0.6 increases the magnitude of pacemaker rate fluctuations, and this effect (of amplitude) is greater at a higher vagal tone.

**Figure 7:**
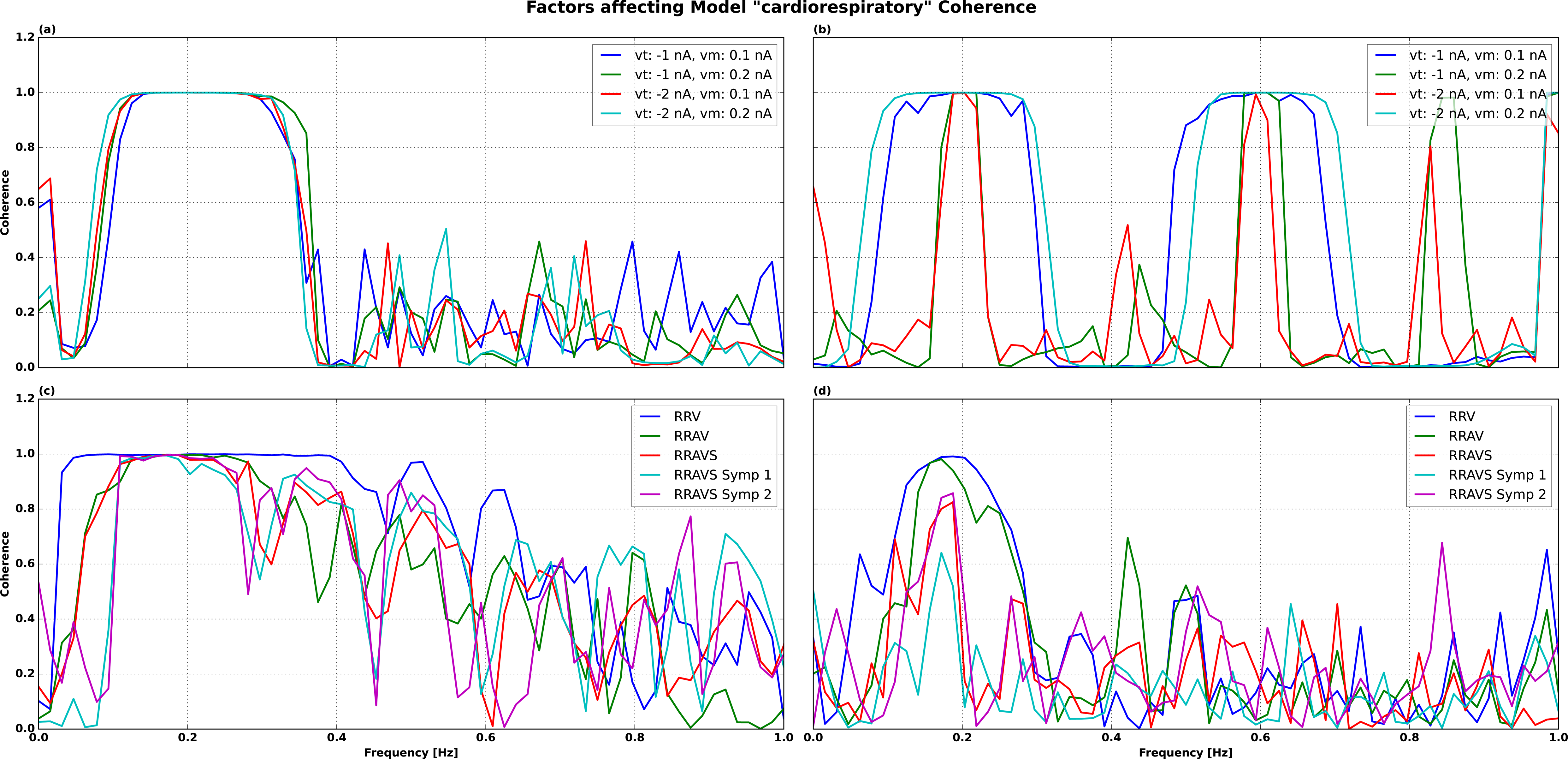
The effect of respiratory variability, sympathetic modulation, autonomic tone and non-linear cardiorespiratory interactions on coherence. Simulated respiratory-vagal input was centred around 0.2 Hz (12 breaths/min). (**A)**-Sinusoidal breathing without logistic transformation of the injected current, (**B)**-Sinusoidal breathing with logistic transformation of the injected current. In both subplots A & B, different values of vago-sympathetic tone (**vt**) and vagal modulation amplitude (**vm**: corresponding to depth of breathing) are simulated, and they reveal no difference in coherence between the simulated conditions, with peak coherence equal to 1. (**C)**-Respiratory variability without logistic transformation. **(D)**-Respiratory variability with logistic transformation. **RRV**-respiratory rate variability, **RRAV**-respiratory rate and amplitude/depth variability, **RRAVS**-respiratory rate and amplitude/depth variability with sighs, **RRAVS Symp 1** - respiratory rate and amplitude/depth variability, sighs, sympathetic modulation, higher sympathetic tone/lower vagal tone (DC-bias=-1.5 nA), **RRAVS Symp 2** - respiratory rate and amplitude/depth variability, sighs, sympathetic modulation, lower sympathetic tone/higher vagal tone (DC-bias=-2 nA). It can be seen that there is greater coherence fall with more respiratory variability, heightened sympathetic modulation/tone and logistic/non-linear transformation of injected current.

#### Simulation of respiratory variability with and without logistic transformation of injected current

With respiratory variability, we show the coherence results without logistic transformation in **Figure 7 (C)** and with logistic transformation in **Figure 7 (D).** Coherence falls progressively with greater respiratory variability (rate & depth variability + sighs: RRAVS in **Figure 7 (D**),) heightened sympathetic tone and modulation (RRAVS Symp 1 in **Figure 7 (**)**D** a)nd logistic transformation of injected current. These results suggest that cardiorespiratory coherence is affected by respiratory variability in general (and sighs in particular), sympathetic modulation (low frequency) and autonomic tone.

### Full-space experiments

The collated plots from all the 96 combinations are shown in **Figures 8-11**. The simulations containing respiratory variability have lower coherence compared to sinusoidal breathing (Figure 8 Vs Figures 9-11). The effect of sinusoidal breathing is shown in **Figure 8**. Pure sinusoidal breathing without sympathetic modulation has a broad-based peak at 0.2 Hz, with peak coherence equal to 1, irrespective of the level of vagal/sympathetic tone. In the presence of sympathetic modulation, the coherence peak is less broad based and begins to fall at lowest vagal tone (AT). Even for the simplest case of sinusoidal breathing, it can be seen that low vagal tone/high sympathetic and sympathetic modulation can cause only a slight decrease in coherence. Without respiratory rate variability, the coherence peak is “un-physiologically” narrow, while normal looking coherence requires the presence of respiratory rate variability. The presence of multiple types of respiratory variability (rate variability, depth variability & sighs) has a greater effect compared to only one type of variability (**Figures 9 Vs Figures 10-11**). For any pattern of breathing & sympathetic modulation, lower the vagal tone (higher the sympathetic tone), lower is the coherence and vice versa. The simulations containing sympathetic modulation have lower peak coherence. The greatest value of respiratory-band cardiorespiratory coherence (centred around 0.2 Hz) is seen with sinusoidal breathing, the highest vagal tone, and no sympathetic modulation (**Figure 8**). The lowest value of peak cardiorespiratory coherence in the respiratory-band (centred around 0.2 Hz) is seen with a conjunction of different types of respiratory variability (rate, depth variability & sighs: respiratory rate variability pattern 2 (rrv2)+ amplitude variability + sighs), lowest vagal tone (highest sympathetic tone) and sympathetic modulation (**Figure 11 (d)**). The results indicate that respiratory irregularity, vagal/sympathetic tone, and sympathetic modulation are the critical factors affecting the value of cardiorespiratory coherence.

**Figure 8:**
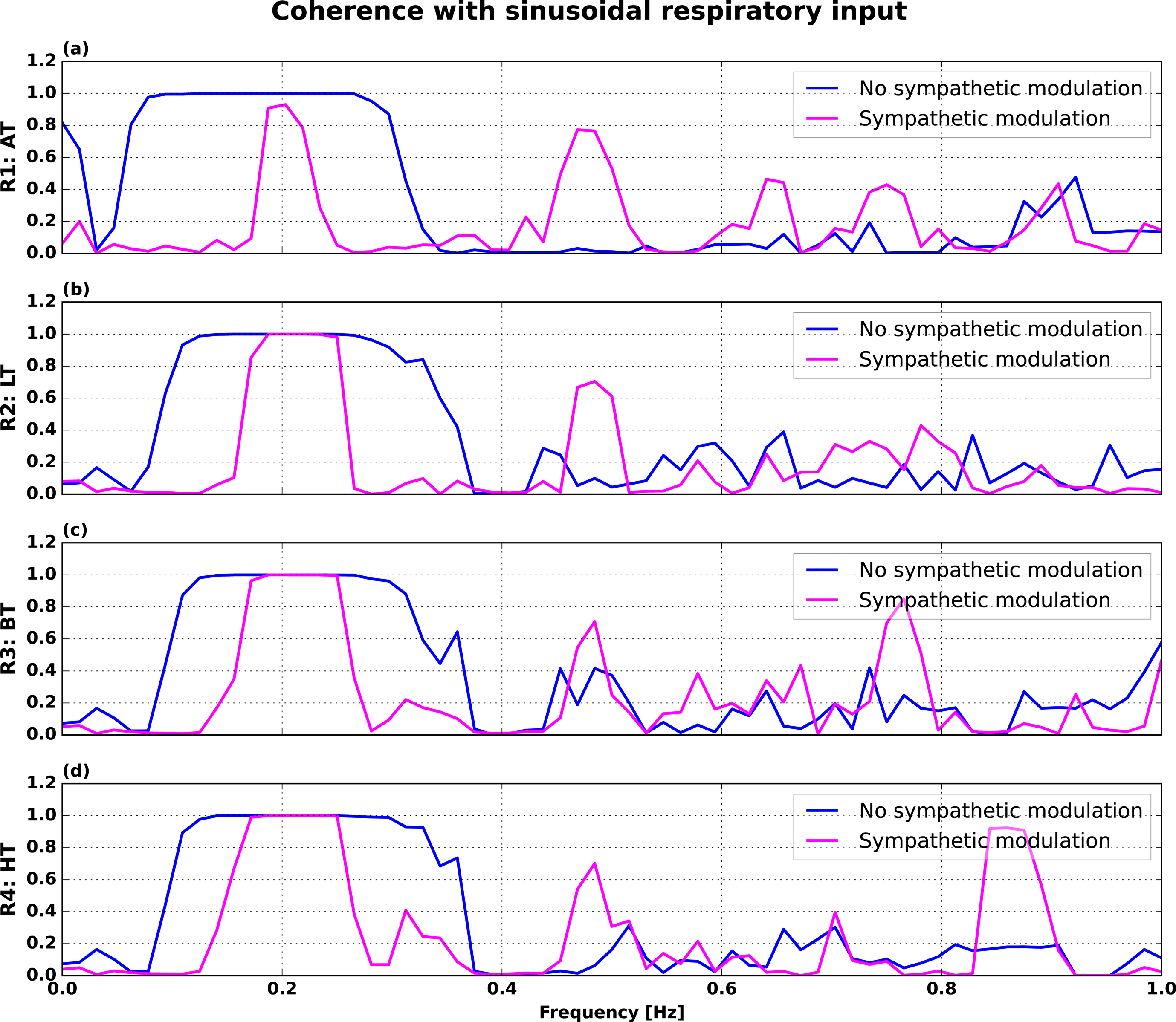
Model Cardiorespiratory coherence in sinusoidal breathing with and without sympathetic modulation. The four rows (R1: → R4:) reflect different levels of autonomic tone, from **AT**, the lowest vagal tone (highest sympathetic tone) to **HT**, high vagal tone (lowest sympathetic tone). The peak coherence begins to fall with highest sympathetic tone (**AT)** and the presence of sympathetic modulation.

**Figure 9:**
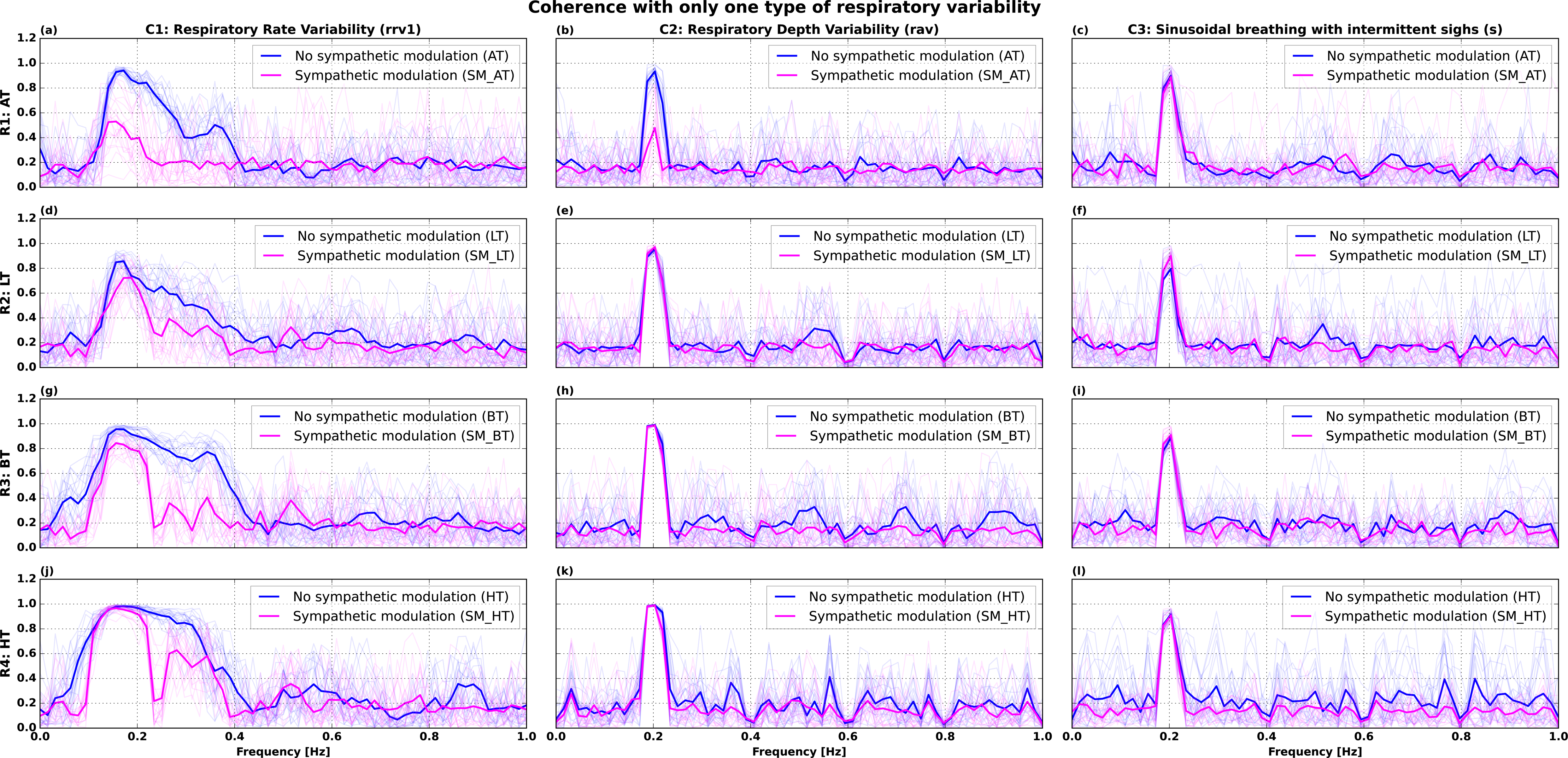
Model coherence at various levels of autonomic tone, one type of respiratory variability. The four rows (**R1: → R4:**) reflect different levels of autonomic tone, from **AT**, the lowest vagal tone (highest sympathetic tone) to **HT**, high vagal tone (lowest sympathetic tone). The three columns (**C1: → C3:**) correspond to different types of respiratory variability viz., rate variability (**rrv1**), depth variability (**rav**) or sighs(**s**). The row (R:) and column (C:) labels of the array of plots define the combination of mechanisms acting on the pacemaker. Respiratory rate variability shown here is Pattern 1 (rrv1) i.e., its frequencies are drawn from power law distribution with exponent parameter=0.3, and it has a mean value of frequency=0.2 Hz. For any type of respiratory variability, the fall of peak coherence (around the respiratory frequency) is greater a lower vagal tone (or higher sympathetic tone)

**Figure 10:**
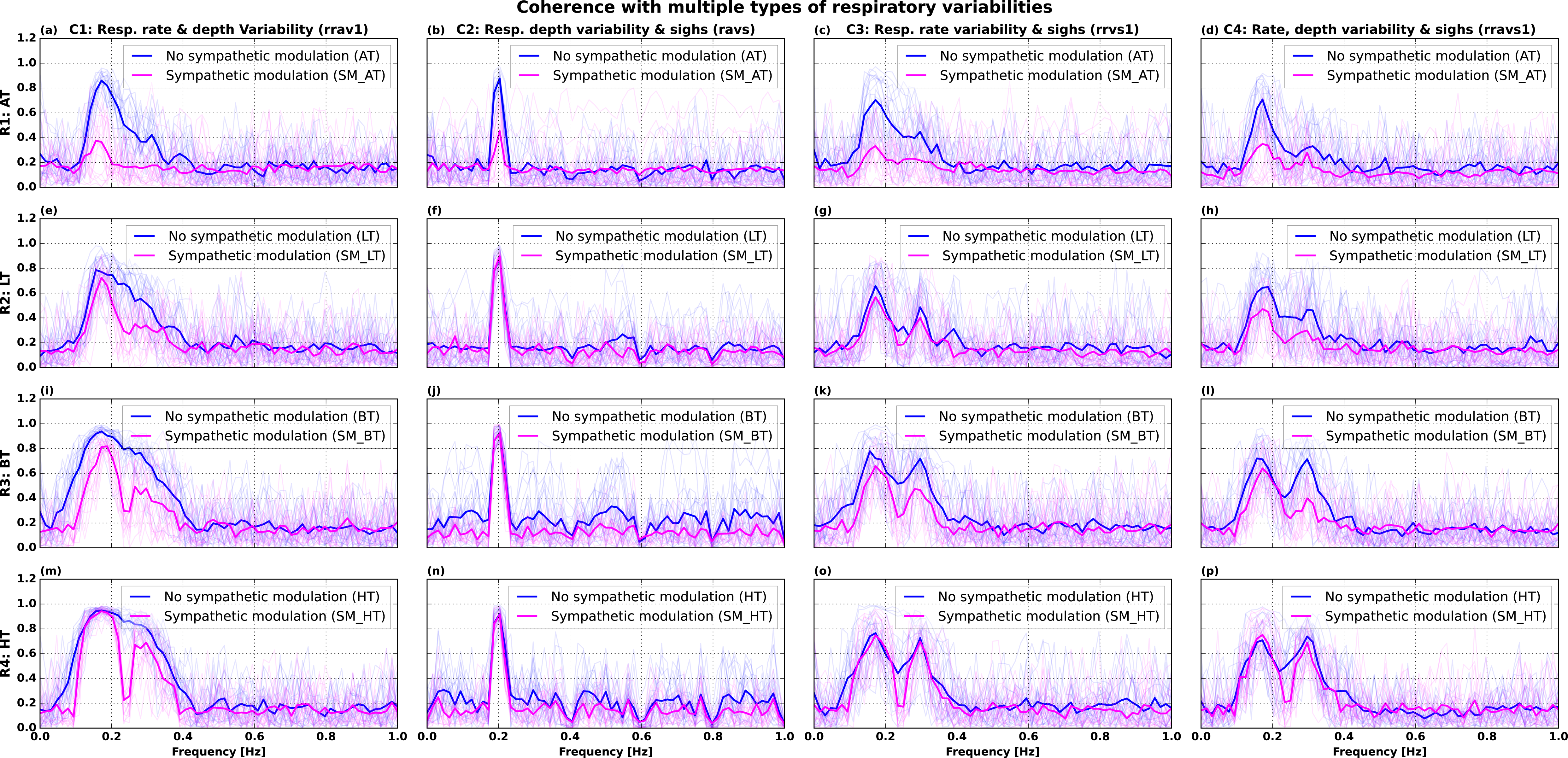
Model coherence at various levels of autonomic tone and respiratory variability. The four rows (**R1: → R4:**) reflect different levels of autonomic tone, from **AT**, the lowest vagal tone (highest sympathetic tone) to **HT**, high vagal tone (lowest sympathetic tone). The four columns (**C1: → C4:**) correspond to different combinations of respiratory variability viz., pattern 1 rate & depth variability (**rrav1**), depth variability & sighs(**ravs**), pattern 1 rate variability & sighs(**rrvs1**) and pattern 1 rate & depth variability & sighs (**rravs1**). Pattern 1 respiratory rate variability has its frequencies drawn from power law distribution with exponent parameter=0.3, and it has a mean value of frequency=0.2 Hz. The fall of coherence is more with greater respiratory variability (of multiple types). This effect is enhanced by lower vagal tone/higher sympathetic tone and presence of sympathetic modulation.

**Figure 11:**
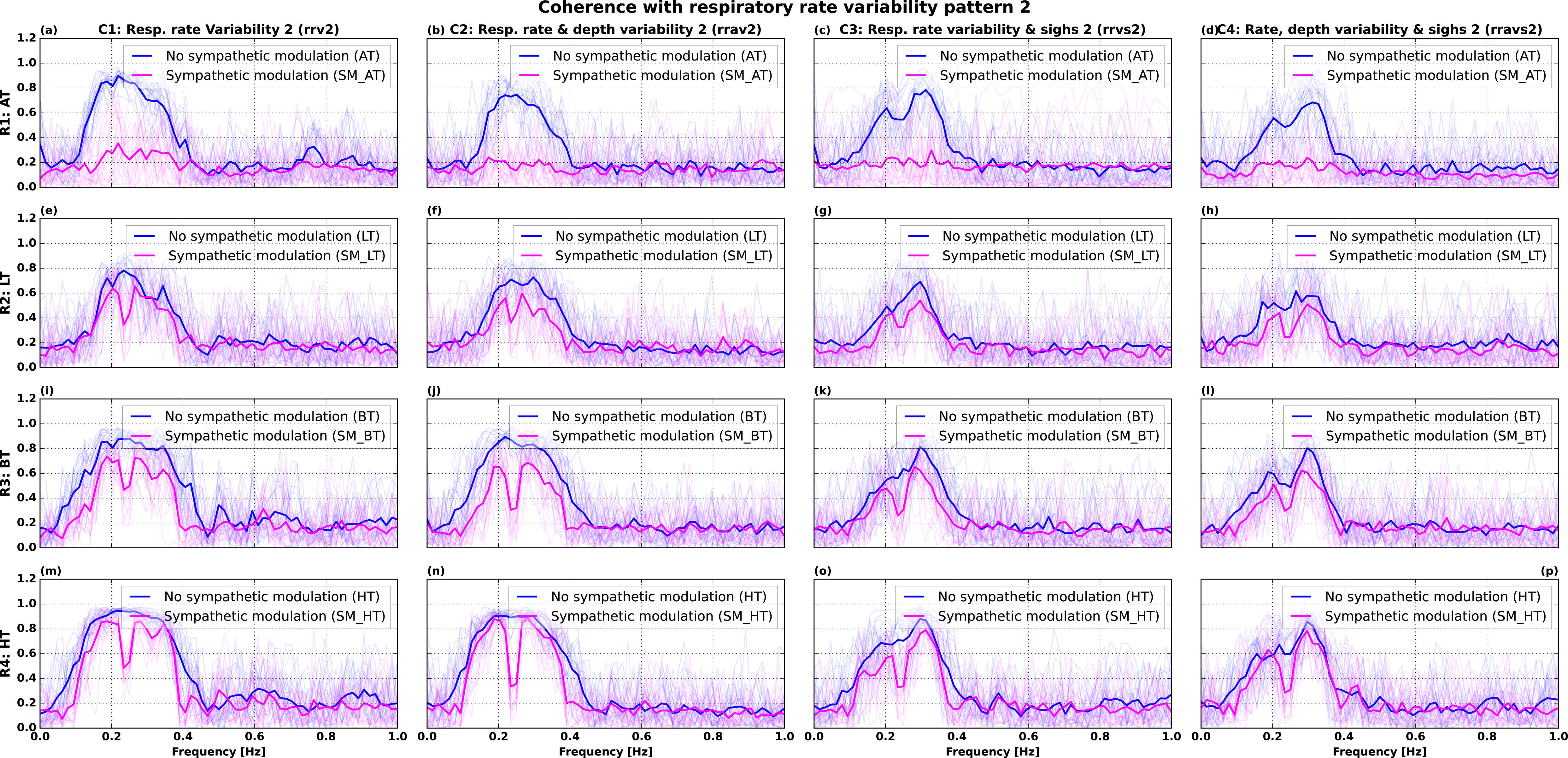
Model coherence at various levels of autonomic tone, respiratory variability, and absence of sympathetic modulation. The four rows (**R1: → R4:**) reflect different levels of autonomic tone, from **AT**, the lowest vagal tone (highest sympathetic tone) to **HT**, high vagal tone (lowest sympathetic tone). The four columns (**C1: → C4:**) correspond to different combinations of pattern 2 respiratory variability viz., rate variability (**rrv2**), rate & depth variability(**rrav2**), rate variability & sighs (**rrvs2**) and rate & depth variability & sighs (**rravs2**). Pattern 2 respiratory rate variability has its frequencies are drawn from power law distribution with exponent parameter=0.8, and it has a mean value of frequency=0.265 Hz. The effect of vagal tone, sympathetic modulation and respiratory variability is qualitatively similar to that in pattern 1 respiration. Least coherence in all the simulations is seen in **rravs2 + lowest vagal tone (AT) +sympathetic modulation.**

In the following, we compare the model derived insights on the factors causing the decline of coherence with the expectation induced cardiorespiratory changes found in the experimental data (CRC study)(11). We show that the respiratory rate & depth variability are similar at rest and anticipation while the sigh frequency increased notably in the anticipatory phase data. The **Figures 12-13**, Appendix **Figures A4-A5** show the distribution of respiratory variability in the participant cohort. **Figure A4** shows the inter-quartile range (IQR) of the respiratory rate in the pre-exercise period for the various types of exercises. **Figure A5** shows the breath-to-breath variability for the respiratory depth. We used IQR as the robust metric to quantify the variability. To quantify sighs, we defined the amplitude criteria as follows: Sighs = amplitude > (75th percentile value of amplitude) + (1.5 × IQR). This definition is based in statistical criterion for an outlier, and is often used in boxplots to identify data outliers. We show that the proportion of sighs (**Figure 12**) to be higher in anticipation of exercise than that in the baseline. **Figure 13** shows a concatenation of baseline and two 5-min pre-exercise respiratory signals in one individual. It can be seen the sigh breaths occur more frequently in the pre-exercise period compared to baseline. Based on the MSE (mean squared error) between experiment-derived (CRC study)(11) average coherence and the model-derived coherence, we extracted the three best-fit models for each of the 11 exercise conditions from 96 model simulations of a combination of mechanisms (**Table 7)**. It can be seen that the presence of a sympathetic modulation (SM) separates pre-exercise conditions from baseline. Most pre-exercise coherence fits better with model simulations involving sympathetic modulation, while baseline has no sympathetic modulation. The highest intensity bilateral bicycle exercise conforms to simulations with the least vagal tone or highest sympathetic tone, sympathetic modulation and respiratory variability. To summarize, we infer the following on the mechanisms for the expectation-induced fall in cardiorespiratory coherence: Pre-exercise state is associated with 1. high sympathetic/low vagal tone and pronounced sympathetic modulation 2. increased respiratory variability in general, and exaggerated frequency of “sighs” in particular.

**Figure 12:**
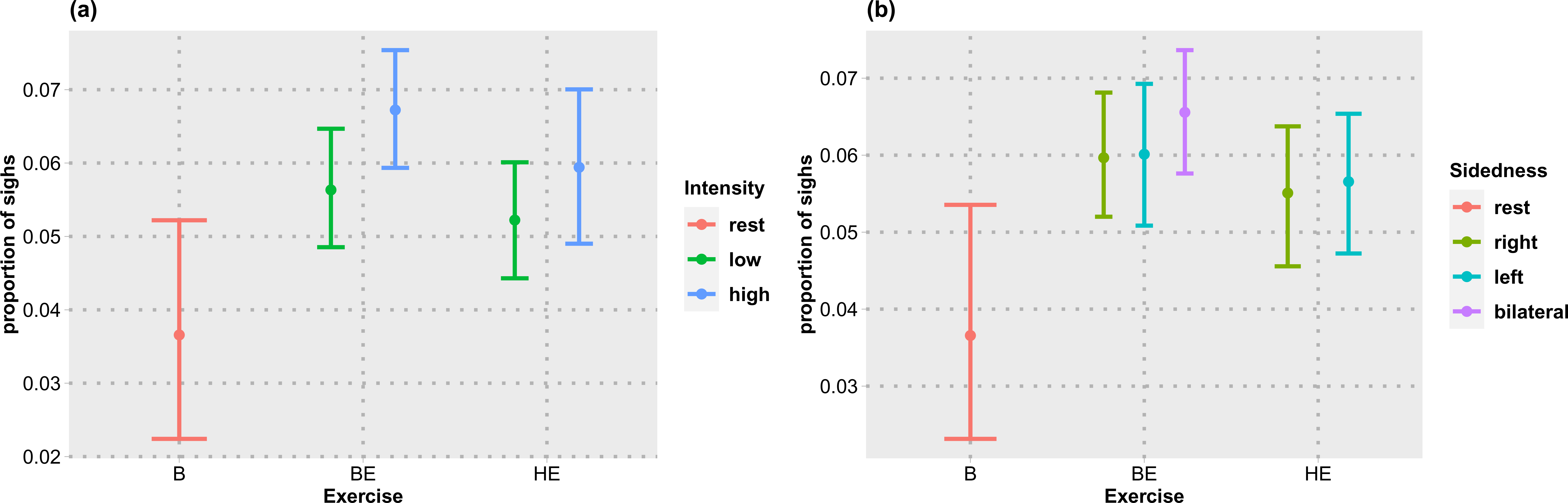
The proportion of sigh breaths from the experimental data. The plots show the average proportion of sighs in the anticipation phase of exercise with the data aggregated by type & intensity or type & sidedness of the exercise. Sigh is defined as a breath whose amplitude exceeds the following cut-off: 75th percentile value for amplitude + 1.5 × IQR of amplitude (statistical outlier deep breath). It can be seen that the frequency of deep sighs increases during anticipation of exercise in load-dependant manner.

**Figure 13:**
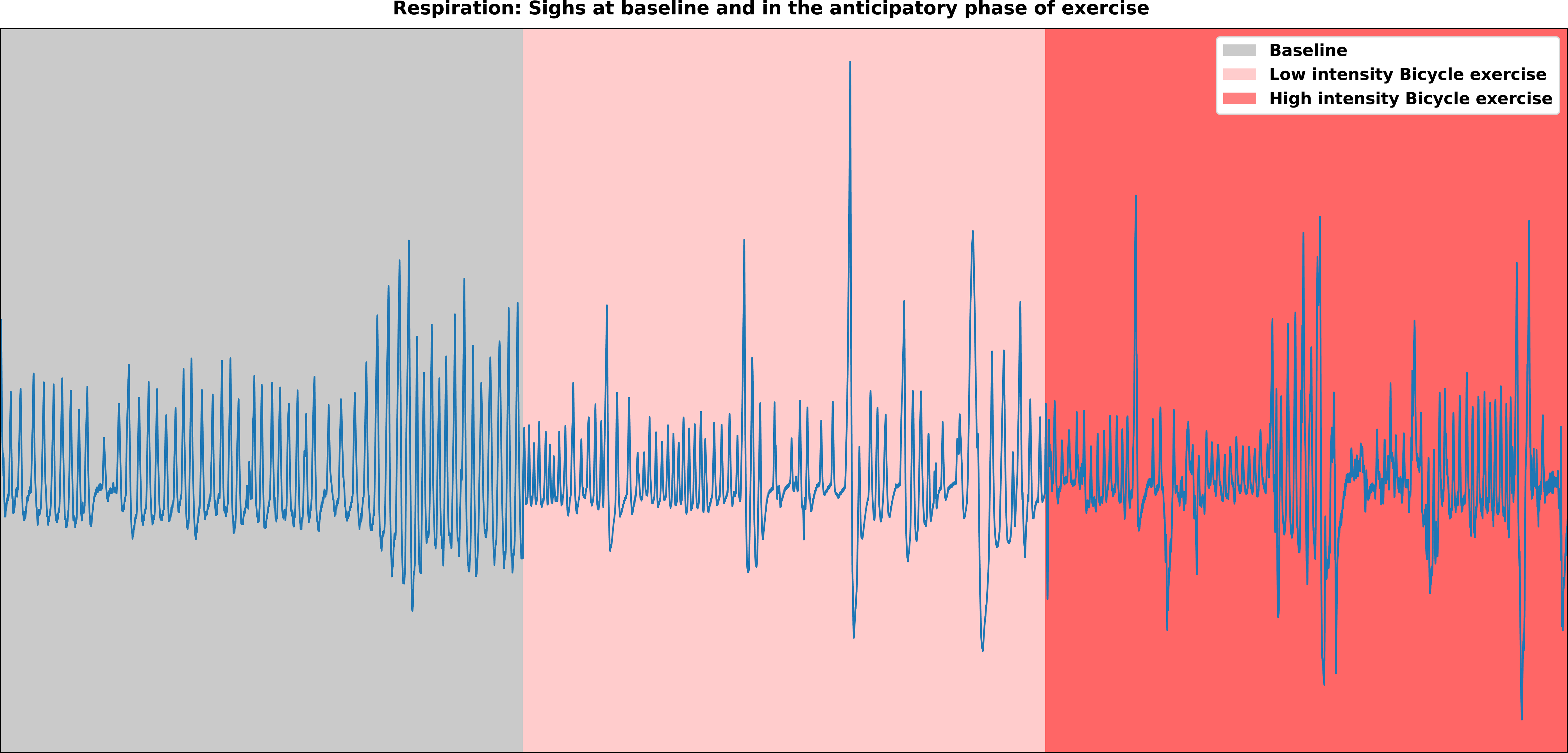
Sigh breaths at rest and during pre-exercise phase. The plot shows three 5-minute stretches of respiration signal at rest (gray background), a single trial of low-intensity bicycle exercise (pink background) & high-intensity bicycle exercise (red background). Large amplitude, sporadically occurring deep breaths are the “sighs”. Sigh breaths can be seen to occur more often in the pre-exercise phase compared to baseline/rest.

## DISCUSSION

We modeled the pacemaker cell of the heart and its modulatory mechanisms to understand the causes of decreased cardiorespiratory coherence induced by the expectation of physical exercise. The modulatory mechanisms evaluated were autonomic tone, vagal/sympathetic modulation, and respiratory variability viz., respiratory rate & depth variability and sighs.

The steady-state firing rate of the pacemaker model is ∼ 105 spikes/minute, which is close to the intrinsic firing rate of the SAN. The heart rate range of a healthy human being at rest is 50-100 beats/minute, which is slightly less than the intrinsic rate. The discrepancy between intrinsic pacemaker rate and heart rate in a healthy human at rest is due to the vagal tone (22,32). Vagal tone is the steady rate of vagal discharge onto the pacemaker cell. The action potential of the model cell shows the various phases, including spontaneous diastolic depolarization or pacemaker potential.

There are several notable differences between the action potential morphology of our model pacemaker cell (**Figure 3**) and the SAN: 1. The upstroke (depolarization) and downstroke (repolarization) speed of the model cell action potential are steeper than that of the SAN. 2. the plateau phase of the SAN action potential is less prominent than model cell 3. The duty cycle of the periodic spikes in the model cell is considerably lesser than that of the SAN. The action potential duration (APD) in the model cell is ∼ 100 milliseconds, while that of SAN > 200 milliseconds. 4. The peak of the action potential in model cell is over +100 mV while it is close to +40 mV in SAN. Therefore, except for the pacemaker rate, model cell action potential morphology differs from that of SAN in almost every other aspect (34,35). We attribute this discrepancy to the fact that only a small subset of ion channels (from a very large number of ion channels that are known to be present in a cardiac cell) was used to design the pacemaker (36). We haven’t included any intracellular calcium storage, release, and uptake mechanisms in the model and this could have affected the action potential properties in the model. Intracellular calcium release/reuptake dynamics play a profound role in the genesis of pacemaker behavior in vivo (25,36–38). Moreover, the ion channels we used in the model were retrieved from a database along with their model parameters. Some of the channels were based on data from animal models, as there is a relative paucity of data from human SAN in this regard. We only iteratively modified the ion channel model parameters to match the pacemaker model cell rate to that of the SAN. For all the above-mentioned reasons, the action potential morphology of model cell is different from that of SAN. This is one of the limitations of the study.

The pacemaker firing rate dropped precipitously from 105 spikes/min as the vagal tone was switched on by injecting an efflux current. The extent of the vagus-induced decline in pacemaker model cell firing positively correlated with the strength of the vagal tone (or the magnitude of negative DC-current) (**Figure 4**). Vagus-induced cardiac deceleration is a physiologically concordant model behavior as the heart rate in healthy humans (22,32,33), as well as animal models(39–41), decreases with an increase in vagal discharge and vice versa. Heart rate at rest is a function of vagal tone. High vagal tone at baseline is seen in trained athletes and it contributes to “sinus bradycardia”in them. High basal vagal tone is thought to confer a number of health benefits, including high exercise capacity, diminished risk of Type 2 Diabetes Mellitus, and all-cause cardiovascular morbidity & mortality, among others(33,42,43). The pacemaker model cell firing rate was steady when the injected current was constant and fluctuated in lock-step with the sinusoidal injected current. This is similar to the behavior of heart rate in response to tonic and phasic vagal discharge (14,44). The rhythmic output from the respiratory central pattern generators is thought be responsible for phasic vagal firing rate fluctuations at the respiratory frequency. The periodic fluctuations of model cell firing rate with sinusoidal input are similar to the RSA i.e., heart rate fluctuations at respiratory frequency(44). The “model RSA” at different levels of vagal tone showed stronger RSA (greater amplitude of model cell firing rate fluctuations) at the higher vagal tone (**Figure 5** & **Table 5**). In healthy humans, heart rate fluctuations have greater amplitude with stronger vagal tone and lower heart rates(45). Magnitude of model cell firing rate fluctuations increased with the amplitude of sinusoidal input. This is similar to large heart rate fluctuations with deep breathing seen in healthy humans. The effect of the amplitude of sinusoidal input interacted with a magnitude of negative DC current. The greater the vagal tone, the greater the effect of sinusoidal amplitude (**Figure 6** & **Table 6**). The physiological counterpart of this model result in humans is a stronger effect of deep breathing on RSA(46,47) at lower heart rates(45). Despite discrepancies in the action potential morphology between model cell & SAN, pacemaker model cell is able to reproduce several phenomena that define the behavior of SAN.

**Table 5:** Effect of vagal tone on pacemaker rate variability.

| S.No | Vagal tone | PR<sub>min</sub> (spikes/min) | PR<sub>max</sub> (spikes/min) | $\Delta_{PR}$ (PR<sub>max</sub>-PR<sub>min</sub>) (spikes/min) |
| --- | --- | --- | --- | --- |
| 1 | Low (-0.3 nA) | 98.23 | 107.64 | 9.41 |
| 2 | Medium (-1 nA) | 87.31 | 100.72 | 13.41 |
| 3 | High (-2 nA) | 66.31 | 81.74 | 15.43 |

The magnitude of steady current injected into the cell to model vagal tone is shown in the parenthesis for each level of vagal tone. The minimum and maximum pacemaker rate in each cycle is averaged over the entire simulation duration. As the vagal tone becomes high, both the maximum (PRmax) and minimum pacemaker rate (PRmin) decrease indicating a fall in the average pacemaker rate with increasing vagal tone. Pacemaker rate variability ( $\Delta_{PR}$ ) is greater at a lower heart rate and higher vagal tone.

**Table 6:**
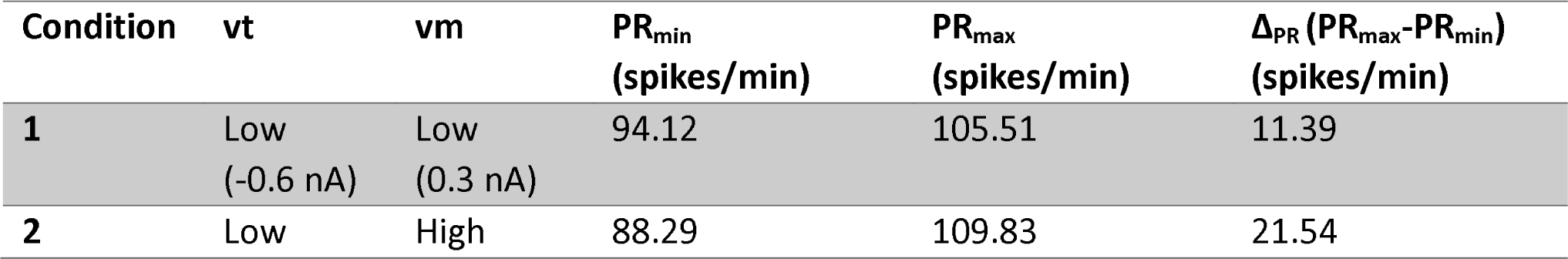
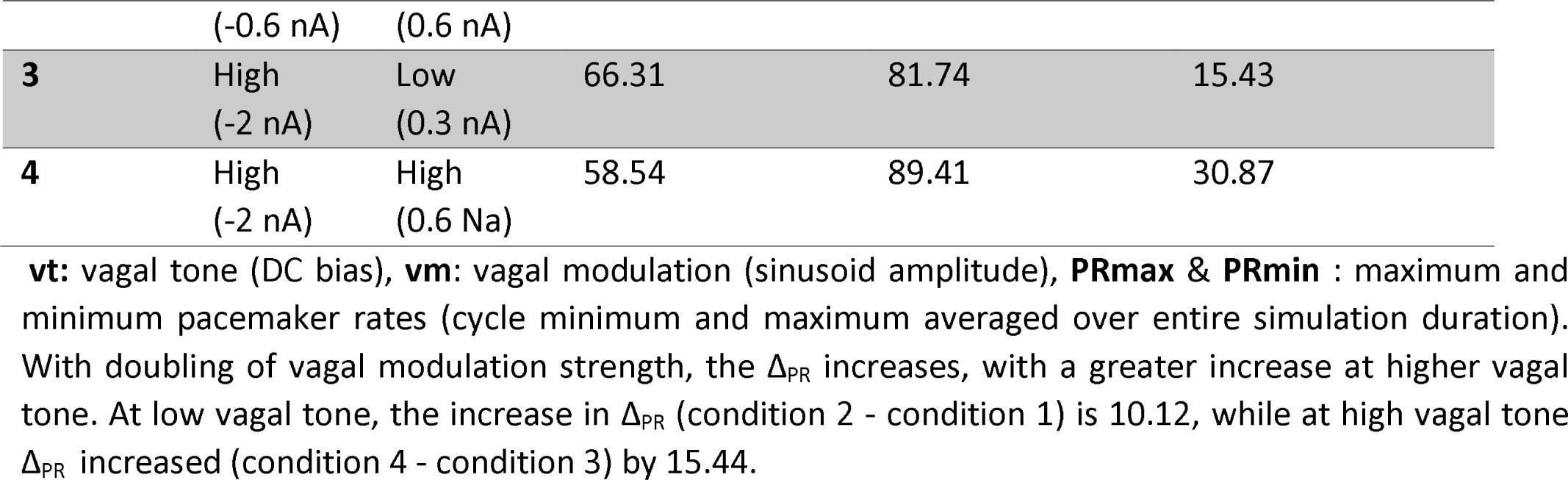
Interaction between the vagal tone and vagal modulation.

We infer the following from our exploration of the effect of various factors on coherence. Change in the autonomic tone alone cannot cause a fall in coherence. An increase in vagal tone increases the time-domain metrics of RSA, like peak-to-peak heart rate variability (**Table 6**) and withdrawal of vagal tone has the opposite effect. However, spectral metrics like coherence are not susceptible to isolated changes in vagal tone without concomitant changes in other factors like respiratory irregularity and non-linear (logistic) transformation of injected current (**Figures 7(A)-(D)**). Coherence decreases as the respiration becomes irregular with increased respiratory rate & depth variability and sighs (**Figure 7(D)**). Coherence falls further as the sympathetic modulation is added and vagal tone is decreased (**Figure 7(D)**). The modeling results implicate respiratory irregularity and a low-frequency sympathetic modulation in the background of vagal tone withdrawal (or sympathetic tone increase) as the putative causal factors for a fall in coherence. The mean squared error between experimental coherence (11 conditions) and model coherence (96 combinations of factors) and evaluation for the best-fit model for each exercise revealed the following. All three best-fit models for the baseline were devoid of sympathetic modulation, while most of the best-fit models for pre-exercise coherence had sympathetic modulation in them. The lowest level of vagal tone (and/or highest level of the sympathetic tone) was associated with high-intensity bilateral bicycle exercise. Respiratory variability/irregularity was present in all the best-fit models for all experimental conditions without exception. The respiratory rate (Appendix **Figure A4**), depth (Appendix **Figure A5**), and sigh (**Figures 12-13**) data from the actual experiment showed that the frequency of sighs differentiated the baseline from pre-exercise conditions. From these results, we conjecture that anticipation of exercise induces alteration in respiratory patterns, including irregular rhythm with intermittent sighs in a load-dependent manner, sympathetic modulation, and vagal tone withdrawal (increased sympathetic tone), causing a fall in cardiorespiratory coherence.

**Table 7:**
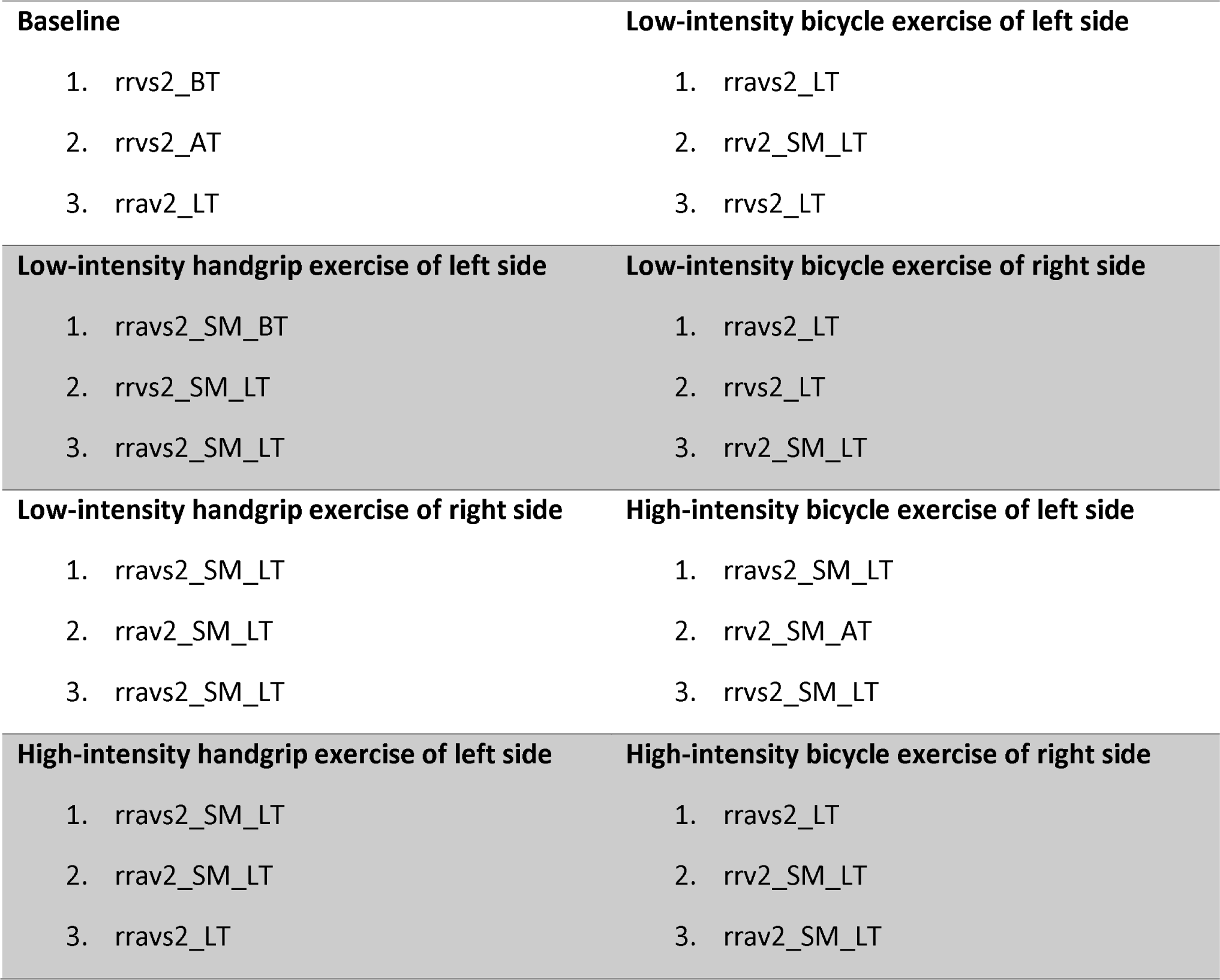
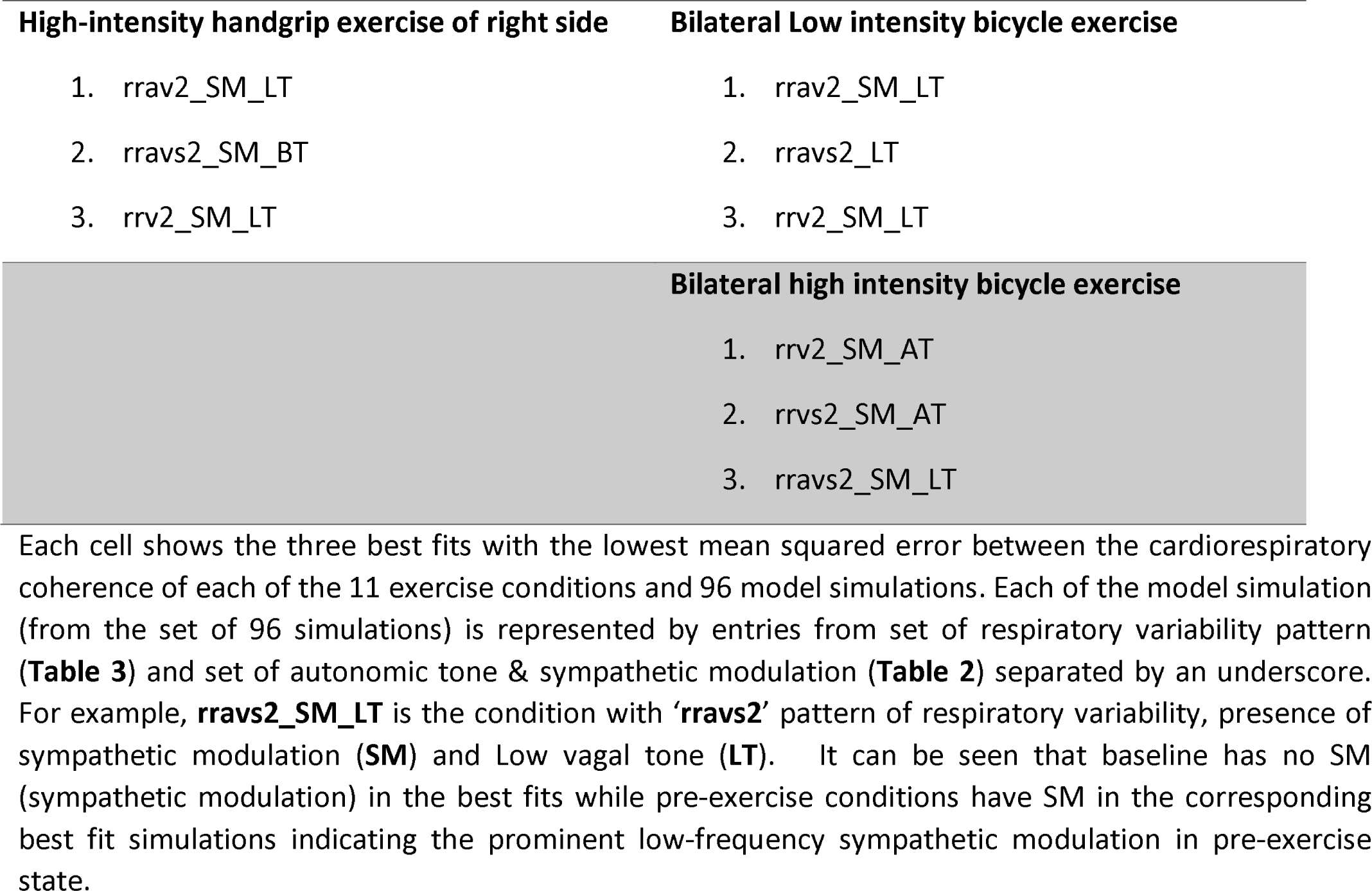
Best fits of coherence for the pre-exercise condition from the 96 model simulations.

There is a substantial body of literature indicating vagal withdrawal and heightened sympathetic response during physical or psychological stress, problem-solving, attentive-arousal states that precede flight-fright-fight response etc.,(3,48–50). Modeling results indicating sympathetic modulation and heightened sympathetic tone during anticipation are consistent with all these studies. Several studies have shown a relationship between stress, panic, arousal etc., with respiratory irregularity (51–53). The frequency of “sighs” increases during stressful situations and emotional states laden with high arousal, anxiety, mental arithmetic, impending public music performances, cognitively intense mental imagery etc (54). It has been suggested that sighs reset the respiratory-cognitive interactions, promote psychophysiological flexibility, facilitate phase transitions between different states, and help adapt to the loads/demands imposed on the physiological systems at the level of the whole organism (55). The causal mechanisms implied by the modeling are consistent with both experimental data and published literature.

The Limitations of our modeling approach are outlined below. The pacemaker model has only a limited subset of ion channels known to be present in the human heart. The ion channels and their parameters are derived from animal models rather than human SAN because of the paucity of human data. The action potential morphology is different from that of human SAN. We have not incorporated any intracellular calcium dynamics-based mechanisms in the genesis of pacemaker activity. Simulation of sympathetic and parasympathetic effects is based on the direct injection of currents bypassing the autonomic nerve spiking and post-ganglionic neurotransmitter kinetics involved in intact organisms. All of these factors constrain the extent of biological realism in the model, and we hope to improve that in our future endeavours. In conclusion, the decline of cardiorespiratory coherence in the expectation of exercise is due to heightened sympathetic tone & modulation, vagal tone withdrawal, respiratory irregularity and frequent “sighing”.

## APPENDIX

### CRC study

The experimental paradigm in the **CRC** study (**C**ardio**R**espiratory **C**oherence in exercise anticipation study) is shown the supplementary **Figure A1.** The study was approved the Institutional ethics committee-Gandhi medical college, Hyderabad (Rc.No.IEC/GMC/2017/XXXVIII). In a cohort of 29 healthy young adult males, single-lead ECG and pneumotrace-abdominal belt plethysmography derived respiratory signal were recorded during the 5-minute anticipation phase and 30 seconds of actual exercise (in real trials). Intensity of bicycle exercise was graded based on the percentage of age-estimate maximal heart rate (Maximum heart rate= 220 – Age (years):: Fox’s formula) achieved during the 30 secs of exercise phase. For the bicycle exercise, low intensity is defined as exercise at which 50% of maximal rate is achieved while high intensity exercise needed cycling to push heart rate to 70% age-estimated maximal heart. Intensity of handgrip exercise was determined by the percentage of maximal isometric handgrip force achieved. Maximum handgrip force was measured; low intensity and high intensity were defined as the handgrip effort required to achieve 35% and 50% respectively of the maximum grip force. Participants were given practice sessions for all exercise types before the experiment commenced. During experiment, they were instructed to look at the screen that provided 1. Timer during the anticipation phase 2. Real-time visual display of exercise intensity i.e., instantaneous heart rate (bicycle exercise) or grip force during the exercise. An auditory cue served to trigger the onset of exercise and the participants were instructed to attain the target intensity at the earliest and sustain it for a total of 30 seconds. Bicycle exercise was done in a left sided, right sided and a bilateral manner while handgrips were done with left hand and right hand in a sequential manner as shown in **Figure A1**. Coherence was computed between ECG derived heart rate and respiratory signal acquitted in the 5-minute anticipation phase. The results are shown in the **Figures A2-A3.** The respiratory period variability and depth variability are shown in **Figure A4** and **Figure A5** respectively.

### Simulation of respiratory variability

We simulated the three types of irregular breathing as follows:

**Respiratory rate variability (rrv)**: Single breath cycles with varying periods were created independently and then appended together to get a signal with cycle-to-cycle period variability. The frequencies for the single breath cycles were randomly sampled from power-law probability distribution using the scipy.stats.powerlaw.rvs() function with the following 2 sets of parameters: (1) **Pattern 1**: rrv1= exponent=0.3, loc= 0.15, scale= 0.25 (2) **Pattern 2**: rrv2= exponent=0.8, loc= 0.15, scale= 0.25. The two rate variability patterns have different mean frequencies: rrv1=0.2 Hz (average breathing rate=12 breaths/min) & rrv2 = 0.265 Hz (average breathing rate=16 breaths/min). At each randomly sampled frequency, a single complete cycle of sinusoidal wave at an amplitude of 1 and initial phase of 3π/2 is created. The initial phase value chosen ensures that each sine-wave cycle morphology is “trough-to-trough” instead of “zero-to-zero” or “crest-to-crest”. The “trough-to-trough” sine wave is akin to a single breath cycle and has 2 parts i.e., initial up-going “trough-to-crest” part, which can be interpreted as inspiration, and the succeeding down going “crest-to-trough” part can be interpreted as expiration. The solitary cycles created above are appended together to create a continuous respiratory signal with frequency/rate variability.

**Respiratory amplitude variability (rav):** To create respiratory amplitude/depth variability without rate variability, we created single sine waves at a constant frequency of 0.2 Hz, but randomly varying amplitude. The random values for the amplitude of individual sine waves are sampled from a Gaussian distribution with mean=1 & standard deviation =0.3. As before, we concatenated single breath cycles at a constant frequency, and randomly fluctuating amplitude into a single signal with pure depth variability.

**Sighs (s)**: Sighs are spontaneous large-amplitude breaths that occur sporadically. Sighs are modeled as follows: From the set of single sine waves, four waves are randomly selected, and their amplitude is set to 3, 4, 6, & 10 times the amplitude of other sine waves. After the exaggeration of select sine waves at random, the sine waves are appended to generate a sigh ridden respiratory signal.

## SUPPLEMENTAL MATERIAL

Supplementary Figures S1-S5; The custom written scripts for simulation, and the relevant data are available in the GitHub repository: https://github.com/AdityaKoppula1985/cardiorespiratory-coherence-exercise-anticipation

## ACKNOWLEDGMENTS

We would like to thank Dr Ram reddy Barra for stimulating discussions on cardiorespiratory coupling and coherence, Dr. GS Prema for encouragement and facilitation of the CRC experimental study.

## GRANTS

This research work was supported by the Department of Science and Technology-Science and Engineering Research Board: SERB/BME/F202/2019-20/G256; Indian Council of Medical Research-Centre of excellence on Medical Devices and Diagnostics: ICMR/BME/F055/2021-22/G402; Department of Science and Technology – Science and Heritage Research Initiative (DST/TDT/SHRI-34/2021).

## DISCLOSURES

Authors have no perceived or potential conflict of interest, financial or otherwise.

## AUTHOR CONTRIBUTIONS

**Aditya Koppula**: Conceived and designed research, performed experiments, analyzed data, interpreted results of experiments, prepared figures, drafted manuscript, edited and revised manuscript; **Kousik Sarathy Sridharan**: Analyzed data, interpreted results of experiments, edited and revised manuscript, approved final version of manuscript; **Mohan Raghavan**: Conceived and designed research, interpreted results of experiments, edited and revised manuscript, approved final version of manuscript.

## Appendix Figure legends

**Figure A1:**
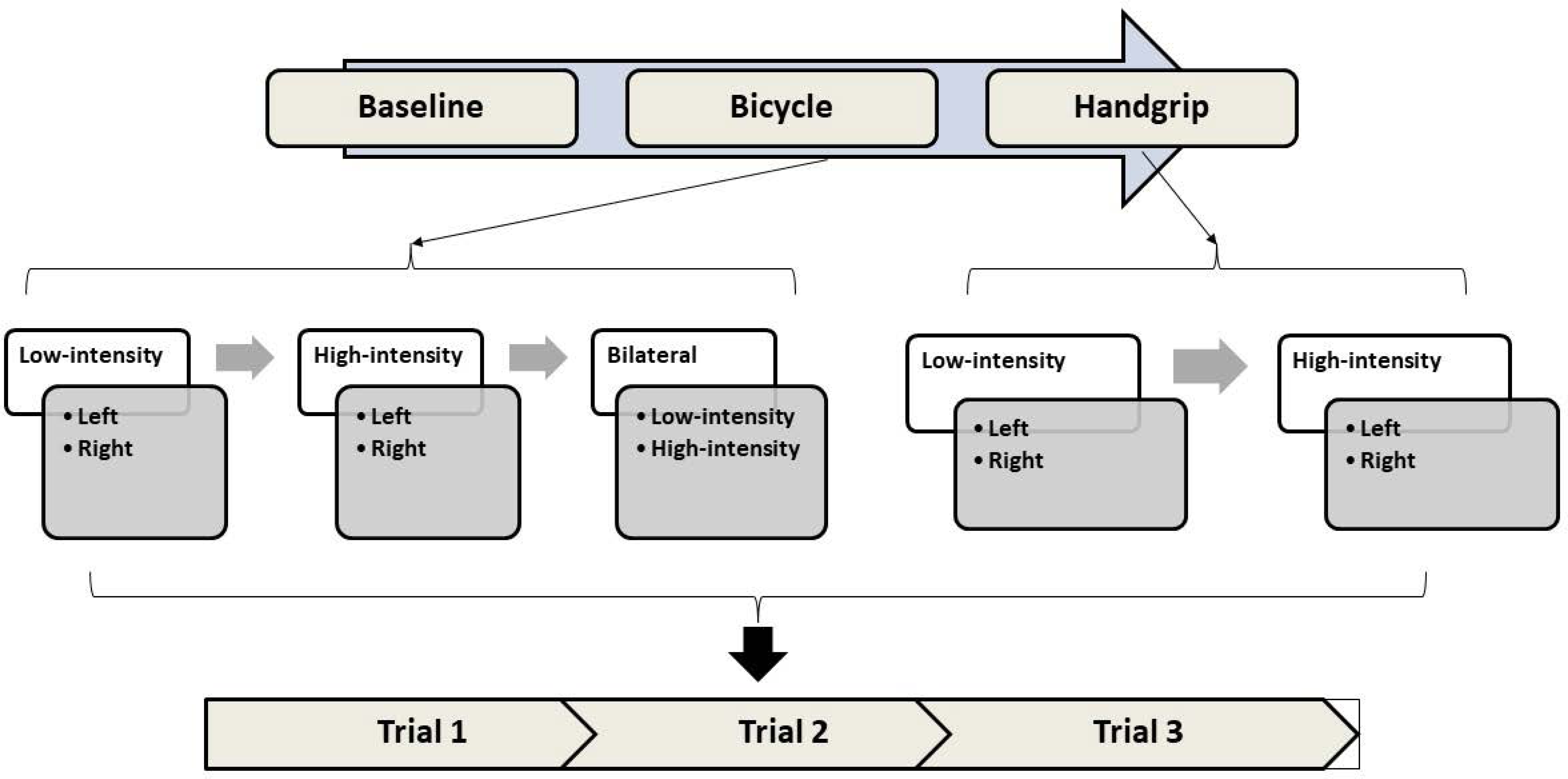
Experimental paradigm in the CRC study. Exercises were performed in the same order for all the participants as shown above. 29 healthy young adult males participated in the experiment consisting of two types of exercise, that is, Bicycling (isotonic) and handgrip (isometric). For both the exercise types, low intensity was followed by high intensity. For each intensity, left sided exercise was performed first followed by the right. Bilateral exercise was performed only for bicycle exercise. For each of the factorial combinations of type, intensity and side of exercise, three repetitions/trials were done. Of the three trials, two were real trials and one was sham trial. The real trials and the sham trials came in a pseudorandom order unknown to the participant. Each real trial had 5-minutes of anticipation phase and 30-seconds of exercise, while sham trials had only the anticipation phase.

**Figure A2:**
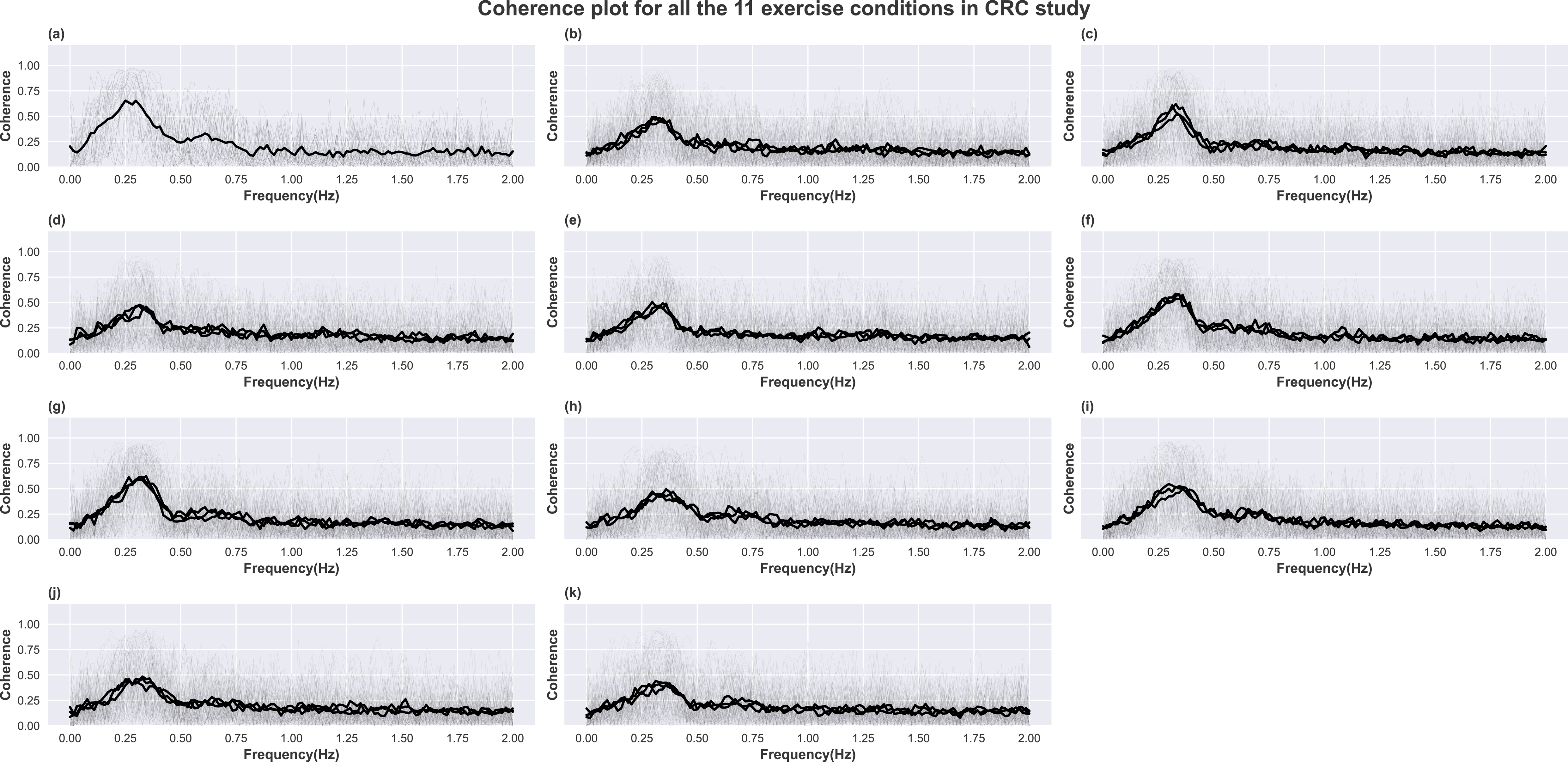
Cardiorespiratory coherence for each exercise condition in the CRC study. Solid curves represent coherence, and the dotted curves represent coherence for surrogate data, created from the same by randomly shuffling the heart rate time series. In health, Coherence plot has a peak in the high frequency region (0.15–0.4 Hz) (3 solid curves, 1 for each trial, in subplots (a–k),that is drastically reduced for shuffled data (3 dotted curves, 1 for each trial, in subplots from (a)–(k). (a) Baseline, (b) handgrip-low intensity-left, (c) handgrip-low intensity-right, (d) handgrip-high intensity-left, (e) handgrip-high intensity-right, (f) bicycle-low intensity-left, (g) bicycle-low intensity-right, (h) bicycle-high intensity-left, (i) bicycle-high intensity-right, (j) bicycle-low intensity-bilateral, (k) bicycle-high intensity-bilateral. Translucent curves in each subplot are the individual coherence curves, whereas the bold-faced curve represents the mean of the same for all the 29 participants. The grand mean coherence for each of the 11 exercise/baseline conditions shown above is termed **“average experimental coherence”** and is compared with **“average model coherence”** to estimate the best fit models for the experimental results.

**Figure A3:**
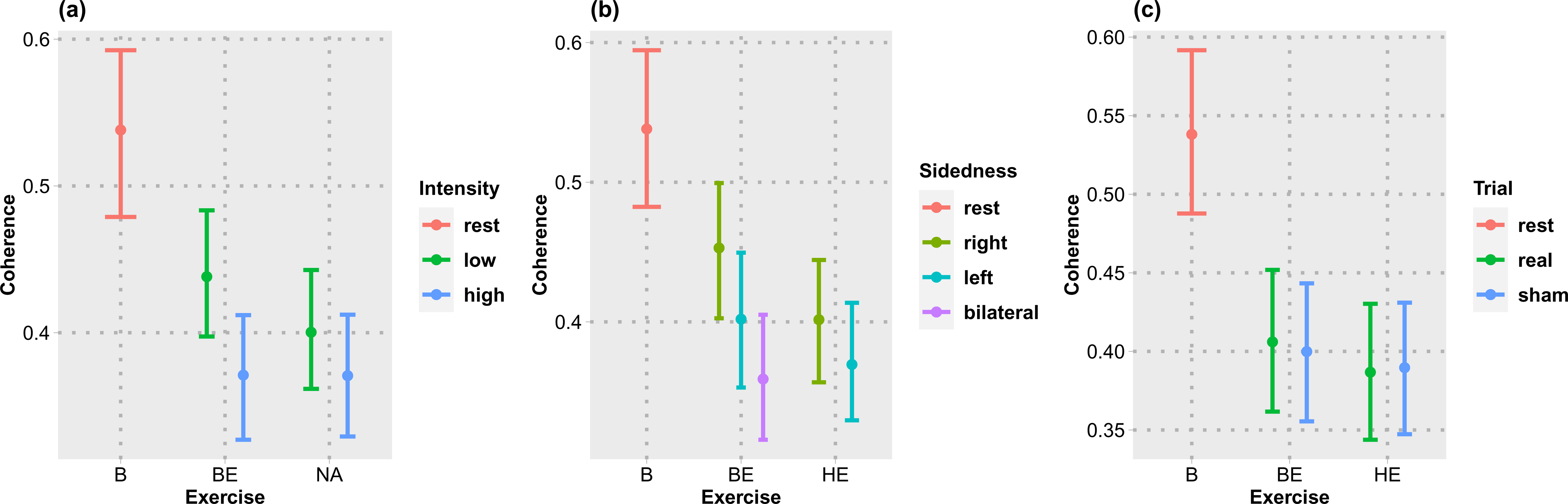
Coherence was averaged over type, intensity, sidedness, and type of trial. Width of the error bars represent 95% confidence intervals estimated using the bootstrap method. (a–c) Coherence changes in each type exercise with the effect of intensity (a), sidedness (b), and trials (c). B, baseline; BE, bicycle exercise; HE, handgrip exercise. The expectation of exercise resulted in a fall of coherence in a load-dependant manner. The coherence decrease was greater for high compared to low intensity exercise, bilateral compared to unilateral exercise. The coherence decreased similarly in both real and sham trials.

**Figure A4:**
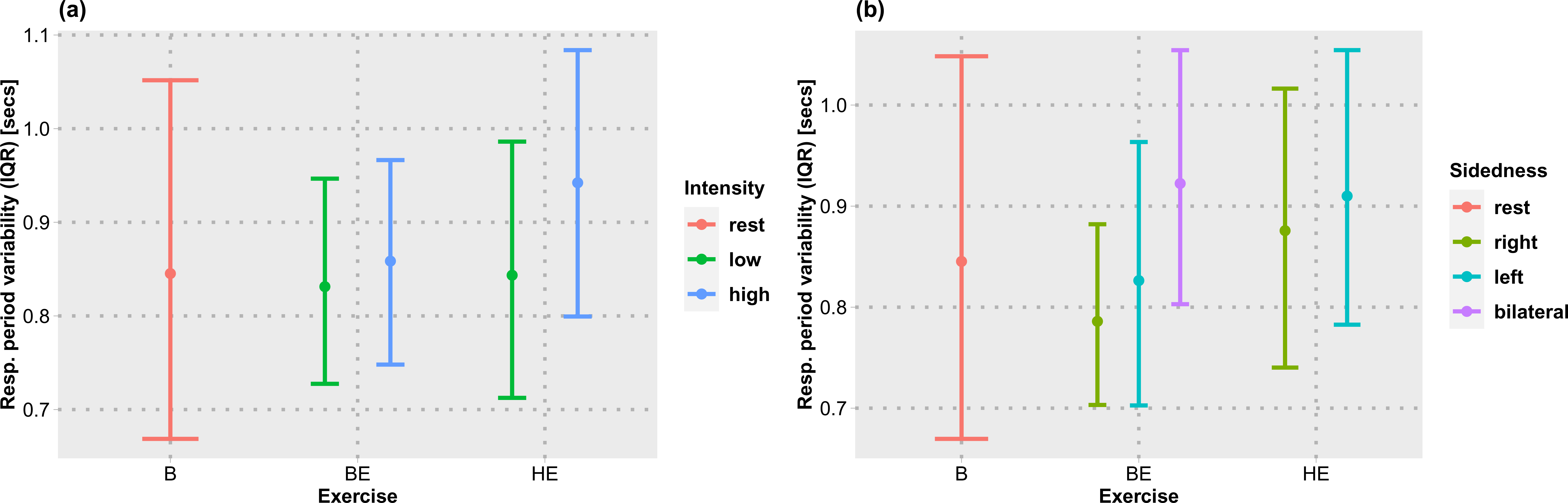
Respiratory period variability from the experimental data. The plots show the average respiratory period variability in the anticipation phase of exercise with the data aggregated by type & intensity or type & sidedness of the exercise. From each 5-min stretch of respiration during the anticipation, the Interquartile range (IQR) of breath-to-breath respiratory time period is computed, averaged, and plotted. In can be seen that there is no difference in the mean respiratory period variability between baseline and pre-exercise conditions.

**Figure A5:**
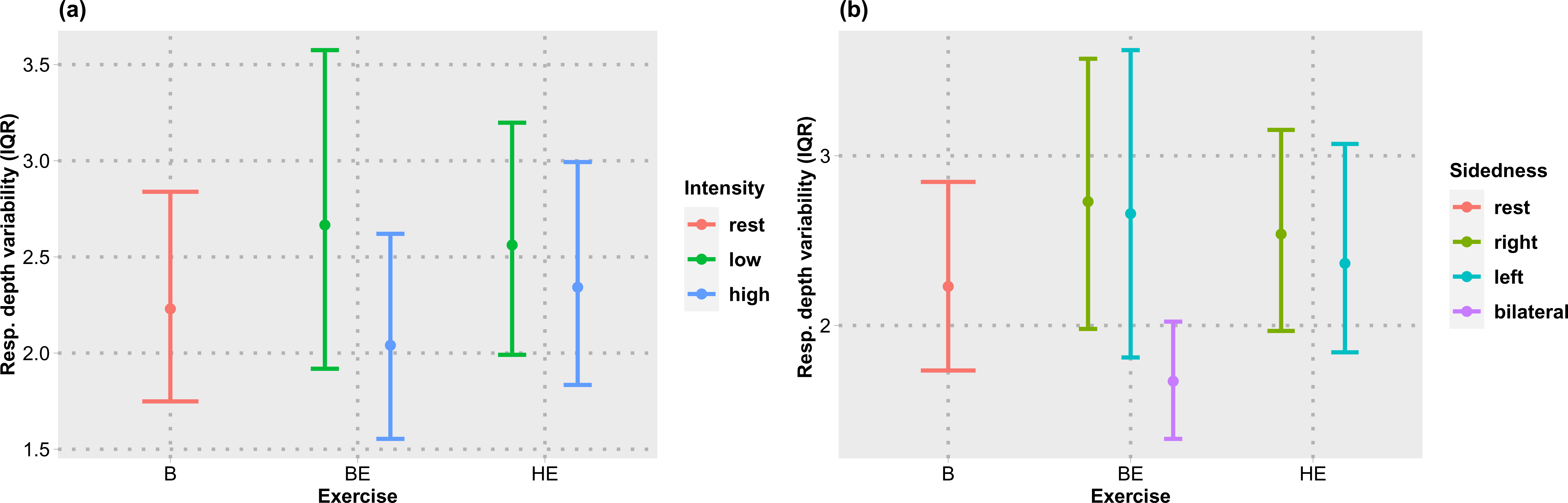
Respiratory depth variability from the experimental data. The plots show the average variability of respiratory depth in the anticipation phase of exercise with the data aggregated by type & intensity or type & sidedness of the exercise. From each 5 minutes of respiration during the anticipation, the Interquartile range (IQR) of the maximum amplitude of each breath is computed, averaged, and plotted. In can be seen that there is no consistent difference in mean respiratory depth variability between baseline and pre-exercise conditions.

## Notes

https://github.com/AdityaKoppula1985/cardiorespiratory-coherence-exercise-anticipation

### Competing Interest Statement

The authors have declared no competing interest.

